# Efficient Estimation of Indirect Effects in Case-Control Studies Using a Unified Likelihood Framework

**DOI:** 10.1101/2021.07.16.452552

**Authors:** Glen A. Satten, Sarah W. Curtis, Claudia Solis-Lemus, Elizabeth J. Leslie, Michael P. Epstein

**Affiliations:** Department of Gynecology and Obstetrics, Emory University, Atlanta, GA; Department of Human Genetics, Emory University, Atlanta, GA; Department of Plant Pathology, Wisconsin Institute for Discovery, University of Wisconsin, Madison, WI

## Abstract

Mediation models are a set of statistical techniques that investigate the mechanisms that produce an observed relationship between an exposure variable and an outcome variable in order to deduce the extent to which the relationship is influenced by intermediate mediator variables. For a case-control study, the most common mediation analysis strategy employs a counterfactual framework that permits estimation of indirect and direct effects on the odds ratio scale for dichotomous outcomes, assuming either binary or continuous mediators. While this framework has become an important tool for mediation analysis, we demonstrate that we can embed this approach in a unified likelihood framework for mediation analysis in case-control studies that leverages more features of the data (in particular, the relationship between exposure and mediator) to improve efficiency of indirect effect estimates. One important feature of our likelihood approach is that it naturally incorporates cases within the exposure-mediator model to improve efficiency. Our approach does not require knowledge of disease prevalence and can model confounders and exposure-mediator interactions, and is straightforward to implement in standard statistical software. We illustrate our approach using both simulated data and real data from a case-control genetic study of lung cancer.

## INTRODUCTION

Mediation techniques^1–4^ are indispensable epidemiological tools for exploring the relationship between an exposure of interest and an outcome variable by investigating how their relationship is affected by intermediate mediator variables. Mediation models partition the total effect that an exposure has on outcome into the indirect effect on outcome explained by the mediator and the direct effect on outcome not explained by the mediator. Classic methods for mediation analysis are embedded within a structural equation modeling (SEM) framework^5–7^ that fit 3 distinct regression models: 1) regression of outcome on exposure, 2) regression of mediator on exposure, and 3) regression of outcome on exposure and mediator. Estimates of direct effect can be based on exposure-outcome relationship in model (3), while estimates of indirect effects are derived from the product of the exposure-mediator relationship in (2) and the mediator-outcome relationship in (3) (or, alternatively, the difference of the exposure-outcome relationship in (1) and the exposure-outcome relationship in (3)). Counterfactual approaches for mediation analysis^8–10^ expanded on the SEM framework to allow for estimates of indirect and direct effects in the presence of potential mediator-exposure interactions as well as nonlinear relationships between variables.

Many studies, particularly in genetic epidemiology, are case-control studies. Mediation analysis for case-control studies often use the influential framework of VanderWeele (VW) and colleagues. The VW framework, published in a pair of landmark papers^10,11^ with over 1100 citations combined, showed how the counterfactual framework could be used to estimate indirect and direct effects on the odds ratio scale for dichotomous outcomes assuming either binary and/or continuous mediators^10,12^.

As described in Materials and Methods, the VW framework conducts inference using two separate regression models: (a) a logistic regression model of disease on exposure and mediator and (b) a separate regression model of mediator regressed on exposure. Estimates and standard errors of total, direct, and indirect effects are then derived based on parameter estimates and standard errors from these two regression models. We note that, while fitting model (a) is straightforward, fitting model (b) is problematic because the sample is non-randomly ascertained under a case-control design, so that parameter estimates based on data from study participants do not necessarily reflect those from the population in general. Studies^13–16^ have found that estimates of the mediator-exposure relationship from a pooled case-control sample are biased when the mediator is associated with disease. If there is no association between exposure and disease, then this bias is generally negligible (unless the mediator is continuous and the effect of mediator on disease is large). However, if there is further association between exposure and disease, then substantial bias can result and be exacerbated both by increasing effect sizes and decreasing disease prevalence.

To circumvent the issue with fitting model(b) in a non-randomly ascertained sample, Valeri and VanderWeele^10^ fit this model using only controls. While this is valid, the exclusion of cases from model (b) means the VW approach loses efficiency relative to methods that could properly model the cases (accounting for ascertainment) when investigating the mediator-exposure relationship. An alternative strategy to incorporate cases into the mediator model was considered by VanderWeele et al.^17^, which employed weighted regression techniques that utilized inverse-probability weighting (IPW) with robust standard errors. A drawback of the IPW approach however is that the method requires knowledge of disease prevalence, which might be difficult to specify, especially when disease prevalence differs between subpopulation groups.

In this article, we propose a likelihood approach for mediation analysis in case-control studies that provides efficiency gains over traditional VW methods. Rather than estimating indirect and direct effects based on two separate regression models, our likelihood approach jointly models the disease, exposure, and mediator together in a unified framework that accounts for correlation in the parameter estimates between the two VW regression models to provide refined estimates of these quantities. Our approach also naturally incorporates cases within the mediator-exposure model and (unlike IPW methods) does not require knowledge of disease prevalence to do so. Our method can also handle confounders and exposure-mediator interactions. The method is simple to implement within the R programming language and we provide code (see Data Availability Statement) for public use. Using simulated data, we show our approach provides more efficient estimates of indirect effects compared to VW approaches (including IPW strategies) for both continuous and binary mediators. We further illustrate our method using genetic and smoking data from a case-control study of lung cancer.

## MATERIALS AND METHODS

### Assumptions and Notation

We assume a case-control study. Define *Y* as disease outcome (1=case, O=control), *A* as an exposure of interest (continuous or categorical), *M* as a possible mediator (continuous or categorical), and *C* as a vector of confounders not influenced by exposure. We wish to fit the mediation model in Figure 1. In particular, we wish to measure the direct effect of *A* on *Y* and the indirect effect of *A* on *Y* carried out through the mediator *M*.

**FIGURE 1:**
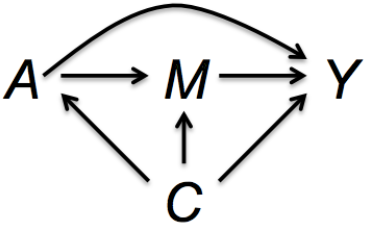
MEDIATION DIAGRAM.

### VanderWeele Framework

The original VanderWeele (VW) technique^10,11^ estimates direct and indirect effects in Figure 1 based on a formal causal inference framework^8,9^. This framework conceptualizes counterfactual and potential outcomes, which define the mechanistic process by which an exposure may causally affect the dependent variable conditional on mediators. Assume a binary exposure and let *Y*(*a*) be the potential outcome for exposure level *a* (*a=0,1*). Then the total effect of *A* on *Y* is defined as *E*[*Y*(1)] - *E*[*Y*(0)], which is the difference between the expectation of *Y* when *A* equals 1 and when *A* equals 0.

We can decompose the total effect (*TE*) of exposure on outcome into the natural indirect effect (*NIE*) and the natural direct effect (*NDE*). To do so, define *M_a_* as the counterfactual value of the mediator *M* if exposure *A* was set to level *a* (*a=0,1*). Next, define *Y*(*a, M_a_*) as the potential outcome if exposure *A* was set to level *a* (*a=0,1*) and mediator *M* was set to level *M_a_*. Then, the counterfactual value of the *NIE* is defined as *NIE* = *E*[*Y*(1, *M*_1_)] – *E*[*Y*(1, *M*_0_)], where *M*_0_ (*M*_1_) is the counterfactual value of the mediator *M* when *A* equals 0 (1). Likewise, *NDE* is defined as *NDE* = *E*[*Y*(1, *M*_0_)] – *E*[*Y*(0, *M*_0_)]. Thus, the total effect equals *TE* = *NIE* + *NDE* = *E*[*Y*(1, *M*_1_)] – *E*[*Y*(0, *M*_0_)]. Additionally, for a given (fixed) value of mediator *M*, we can define the controlled direct effect (*CDE*) as *CDE* = *E*[*Y*(1, *M*)] – *E*[*Y*(0, *M*)], which can change for different values of *M* (as we will show). The framework can estimate direct and indirect effects despite only one of the potential outcomes being observable for a given subject, provided four assumptions hold: (I) no unmeasured exposure-outcome confounding, (II) no unmeasured mediator-outcome confounding, (III) no unmeasured exposure-mediator confounding, and (IV) no mediator-outcome confounder affected by exposure.

For case-control studies with a binary outcome, the VW techniques^11^ extended the definitions of *NIE, NDE*, and *CDE* to the odds ratio scale. On this scale, the total effect (conditional on confounders *C*) is defined as

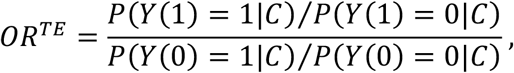

where *P*(*Y*(*a*) = 1| *C*) is the probability of disease when *A=a. TE* can be partitioned into the product of the *NIE* and *NDE*, which are defined as

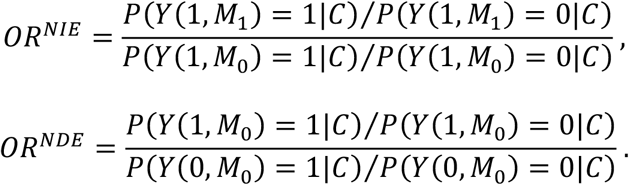

The VW framework estimates *OR^TE^, OR^NDE^, OR^NIE^* by first fitting regression models to the observed data. To model the disease data, the VW framework applies the logistic model

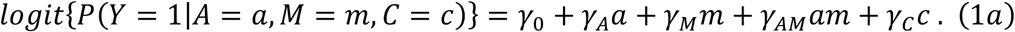

To model the mediator (continuous or binary), VW uses one of the following regression models

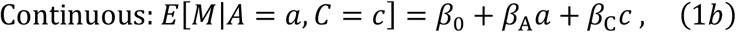

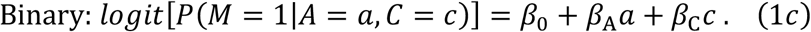

For a continuous mediator, the VW framework estimates *NIE, NDE*, and *CDE* on disease outcome going from exposure level *a\** to *a* (assuming assumptions I-IV above hold) under a rare disease assumption (i.e. disease prevalence approximately below 5%) as

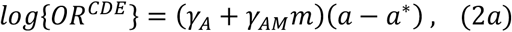

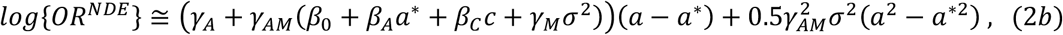

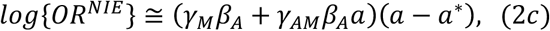

where *σ*^2^ in equation (2b) is the residual variance from fitting model (1b), which is assumed to be normally distributed. Note that *CDE* in (2a) depends on level of the mediator (as previously mentioned). In contrast, for a binary mediator, the VW framework estimates *NIE*, *NDE*, and *CDE* on disease outcome going from exposure level *a\** to *a* under a rare disease assumption as

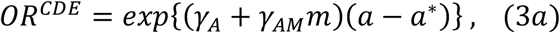

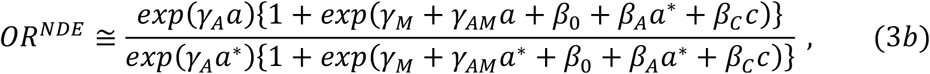

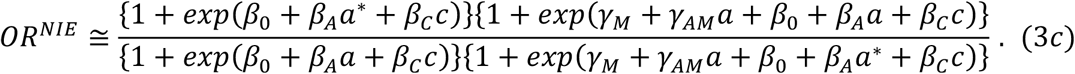

For either a continuous or binary mediator, the framework then estimates the *TE* on disease outcome going from exposure level *a\** to *a* as *OR^TE^ = OR^NDE^OR^NIE^*. Once we estimate these quantities, we can derive the standard errors of *OR^TE^, OR^NDE^, OR^NIE^* on the log scale by applying the delta method using the formulae outlined in the Appendix of Valeri and VanderWeele^10^ and presented here in Supplementary Materials. We can then perform hypothesis testing of *OR^TE^, OR^NDE^, OR^NIE^* using Wald tests.

The VW framework’s estimation of *NDE, NIE*, and *CDE* for both continuous mediators in (2) and binary mediators in (3) requires parameter estimates from the mediator models in (1b) and (1c). Fitting models (1b) and (1c) is difficult because the sample is ascertained based on disease and does not represent a random sample from a population as assumed by *E*[*M*|*a*, *c*] in (1b) and *logit*[*P*(*M* = 1|*a, c*)] in (1c). To circumvent this complication, the VW framework utilizes the rare-disease assumption to justify fitting *E*[*M*|*a*, *c*] and *logit*[*P*(*M* = 1|*a, c*)] in (1b) and (1c) using control data only. Alternatively, as in VanderWeele et al.^17^, one can instead incorporate cases into the mediator models in (1b) and (1c) using IPW such that cases are utilized in the model but their contributions (compared to controls) are downweighted relative to their sampling proportion. For IPW regression, the standard errors of the coefficients in (1b) and (1c) are then derived using robust Huber-White procedures^18,19^.

### Unified Likelihood Approach

Here, we propose a unified likelihood approach that jointly models disease, exposure, and mediator data together while incorporating cases into the mediator model accounting for ascertainment, using the rare disease approximation. Letting *j* index subject and assuming all subjects are independent, we initially define the joint (prospective) likelihood as 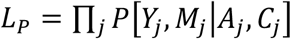, which we then factor into

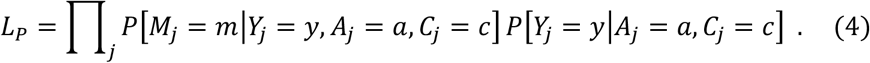

We note that we could replace *P*[*Y_j_* = *y*|*A_j_*, = *a*, *C_j_* = *c*] in (4) with *P*[*A_j_* = *a*|*Y_j_* = *y*, *C_j_* = *c*] and instead perform inference using the retrospective likelihood *L_R_* = Π*_j_ P*[*A_j_*, *M_j_*|*Y_j_*, *C_j_*]. However, the prospective likelihood is often preferred to the retrospective likelihood as *Y*|*A,C* in the former likelihood is easier to model (e.g. using logistic regression) than *A*|*Y,C* in the latter likelihood; standard results^20^ assure that *L_R_* and *L_P_* are proportional if a nonparametric distribution is chosen for *A|C*, and that only the intercept is affected.

We first describe how to model *P*[*M_j_* = *m|Y_j_* = *y, A_j_* = *a, C_j_* = *c*] in (4). For controls, we assume a rare disease such that *E*[*M_j_* = *m|Y_j_* = 0, *A_j_* = *a, C_j_* = *c*] ≅ *β*_0_ + *β_A_ a* + *β_C_ c* (for a continuous mediator) or *logit*[*P*(*M* = 1|*Y_j_* = 0, *A_j_* = *a, C_j_* = *c*)] ≅ *β*_0_ + *β_A_a* + *β_C_c* (for a binary mediator). These choices are the same as usually used when using the VW framework.

Data from cases can be incorporated into the mediation model; these contributions are discarded in the VW framework. To accomplish this, note that we can write *P*[*M_j_* = *m*|*Y_j_* = 1, *A_j_* = *a,C_j_* = *c*] as^21,22^;

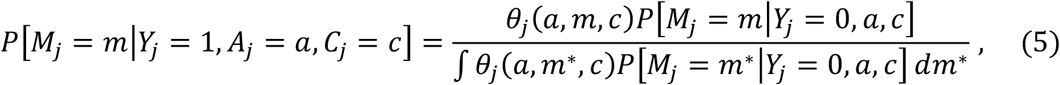

where

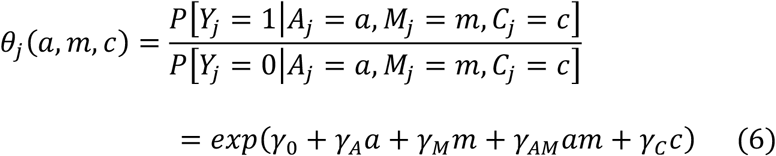

is the disease odds given exposure, mediator, and confounders (allowing for possible mediator-exposure interaction, if desired). Note that *θ_j_* (*a, m, c*) is the same quantity fit in equation (1a) in our summary of the VW framework.

Equation (5) shows that we can naturally model the mediator in the cases (accounting for ascertainment) as a function of the disease odds in (6) and the mediator model in the controls, without requiring external information like disease prevalence (which IPW methods require). The denominator in (5) corresponds to the disease odds given exposure and confounders^21,22^. That is,

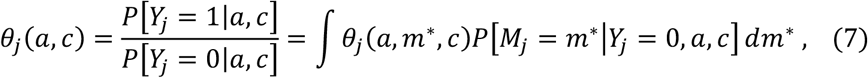

which becomes

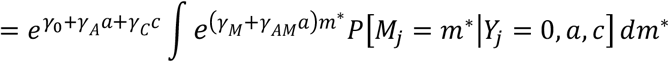

if the model in equation (6) is used. *θ_j_* (*a, c*) in (7) has a closed form whenever the moment generating function for *P*[*M_j_* = *m*\* |*Y_j_* = 0, *a*, *c*] has a simple or closed form, which simplifies inference. For example, if the mediator *M* is normally distributed in controls with mean *E*[*M_j_* = *m*|*A_j_* = *a, C_j_* = *c, Y_j_* = 0] = *β*_0_ + *β_A_a* + *β_C_c* and variance *σ*^2^, then we show in Supplementary Materials that *θ_j_*(*a, c*) = exp(*γ*_0_ + *γ_A_a* + (*γ_M_* + *γ_AM_a*)(*β*_0_ + *β_A_a* + *β_C_c*) + 0.5(*γ_M_* + *γ_AM_a*)^2^*σ*^2^ + *γ_C_c*). For a binary *M*, we simply replace the integral in equation (7) with a sum over the two levels of the mediator.

To complete construction of *L_P_* in (4), we model *P*[*Y_j_* = *y*|*A_j_* = *a, C_j_* = *c*] using a logistic regression model based on *θ_j_*(*a, c*) in (7), such that 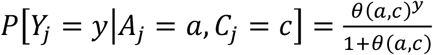 with the assurance that the only difference between using *L_P_* and *L_R_* for case-control sampling is in the intercept^20^ *γ*_0_. Combining all the pieces, we can simplify our likelihood for inference in (4) to give

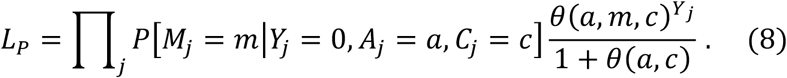

We propose to base inference on *L_P_* in (8); parameter estimates can be obtained by maximizing (8) with respect to (*γ*_0_, *γ_A_*, *γ_M_*, *γ_AM_*, *γ_C_*, *β*_0_ *β_A_*, *β_C_*, *σ*^2^) using standard optimization algorithms like Quasi-Newton, and the covariance matrix of the estimated parameters can be obtained either from the information matrix or a ‘sandwich’ variance estimator. We can then substitute these estimates into equations (*2a*-*2c*) or (*3a–3c*) to obtain estimates of *OR^TE^, OR^NDE^, OR^NIE^*, and *OR^CDE^* and use the delta method results shown in Supplementary Materials (based on the Appendix of Valeri and VanderWeele^10^) to obtain standard errors and construct confidence intervals of these quantities. We can perform hypothesis testing of the parameters or measures of direct and indirect effects using Wald statistics based on the estimates and standard errors.

### Simulation Studies

We performed simulation studies to evaluate the efficiency gains of our likelihood for mediation analysis relative to the original VW method. We assumed a continuous exposure *A* that followed a standard normal distribution and a binary covariate *C* that followed a Bernoulli(0.5) distribution. Using *A* and *C*, we generated a continuous mediator *M* using equation (1b) or generated a binary mediator *M* using equation (1c) using parameter values shown in Table 1. We then used *M, A, C* to generate disease status using equation (1a) and parameter values shown in Table 1, assuming a value for the intercept that yielded a population prevalence of 0.03. As shown in Table 1, we considered models that explicitly assumed an exposure-mediator interaction as well as models that assumed no interactions. We chose parameter values to yield models with *log*(*OR^TE^*) = 0.145, *log*(*OR^NDE^*) = 0.1, and *log*(*OR^NlE^*) = 0.045 for both a continuous and binary mediator, with and without a mediator-exposure interaction effect. For a specific model, we prospectively generated subjects until we obtained 300 cases and 300 controls. For all simulation settings, we generated 1000 replicates per setting.

**TABLE 1:** SIMULATION MODELS.

|  | Continuous Mediator |  | Binary Mediator |  |
| --- | --- | --- | --- | --- |
|  | No Interaction | Interaction | No Interaction | Interaction |
| $\gamma_0$ | -3.476 | -3.476 | -3.476 | -3.476 |
| $\gamma_A$ | 0.100 | 0.077 | 0.100 | 0.049 |
| $\gamma_M$ | 0.300 | 0.200 | 0.420 | 0.320 |
| $\gamma_{AM}$ | X | 0.100 | X | 0.100 |
| $\gamma_C$ | 0.050 | 0.050 | 0.100 | 0.100 |
| $\beta_0$ | 0.100 | 0.100 | -0.310 | -0.310 |
| $\beta_A$ | 0.150 | 0.200 | 0.430 | 0.430 |
| $\beta_C$ | 0.050 | 0.050 | 0.100 | 0.100 |
| $\sigma^2$ | 0.500 | 0.500 | X | X |
| $\log(OR^{TE})$ | 0.145 | 0.145 | 0.145 | 0.145 |
| $\log(OR^{NDE})$ | 0.100 | 0.100 | 0.100 | 0.100 |
| $\log(OR^{NIE})$ | 0.045 | 0.045 | 0.045 | 0.045 |
| $\log(OR^{CDE})$ | 0.100 | 0.077 | 0.100 | 0.049 |

$OR^{TE}$ , $OR^{NDE}$ , $OR^{NIE}$ , and $OR^{CDE}$ calculated where $a^* = 0$ , $a = 1$ , $c = 0$ , and $m = 0$

For each simulation analysis, we compared the performance of our likelihood approach both to the original VW method that discards cases within the mediator models of (1b) and (1c) as well as the IPW-version of VW that incorporates cases into the mediator models of (1b) and (1c) using inverse probability weighting. For IPW analyses, we derived weights using the same strategy of VanderWeele and Vansteelandt^11^. Letting π denote disease prevalence and *p* denote proportion of cases in the case-control sample, we weighted each case by 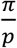 and each control by 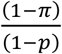). For all IPW simulations, we assumed the disease prevalence of the weighting was correctly specified.

All analyses were conducted in the R programming language. We fit our likelihood to simulated datasets using a quasi-Newton algorithm implemented in the R ‘optim’ function. We fit the VW methods using the R functions ‘glm’ and ‘lm’. For the IPW method, we constructed robust Huber-White standard errors of the mediator model using the ‘vcovHC’ function in the R library *‘sandwich’*.

### Application to Case-Control Genetic Study of Lung Cancer

Lung cancer is the leading cause of cancer-related deaths across the world^23^ and represents a major public health concern. A well-known environmental risk factor for lung cancer is tobacco smoking, with long-term smokers having a 10-fold increased risk of lung cancer relative to non-smokers^24^. Beyond smoking and other environmental risk factors, genetics are also known to broadly play a role in lung-cancer risk^25,26^ with several specific risk loci being identified within genome-wide association studies (GWAS) of the disease ^27,28^.

One established risk loci of lung cancer identified by GWAS resides on chromosome 15q25.1^29^. The associated region contains nicotinic acetylcholine receptor subunit genes that encode proteins that form receptors that bind nicotine. A number of studies suggest that this risk locus is also associated with measures of smoking behavior^30–33^. Given the known relationship between smoking behavior and lung cancer, it is worth inquiring whether the SNPs in this locus influence lung cancer risk partially or completely through smoking behavior. To explore these hypotheses, we utilized genetic, smoking pack-year, and disease data from the GENEVA GWAS of Lung Cancer and Smoking^34,35^ available through dbGaP (accession number phs000093.v2.p2; see Data Availability Statement). The GENEVA GWAS dataset consists of 2695 cases and 2779 controls genotyped for 508,916 post-QC SNPs across the genome.

We first used the GENEVA data to perform a genome-wide analysis of the post-QC autosomal SNPs with lung cancer using logistic regression, adjusted for gender and the top three principal components of ancestry (Manhattan and quantile-quantile plots are shown in Figures S1 and S2, respectively). For each SNP, we assumed an additive model and coded the genetic predictor as the number of copies of the minor allele that a subject possessed. Within the chromosome 15q25.1 locus, we identified strong associations of several SNPs with lung cancer in GENEVA with SNP *rs12914385* (*p=1.887×10^−15^*) yielding the top signal in the region (see Figure S3). We next conducted a GWAS of a dichotomized version of smoking pack-years (split at the median value of 3 pack-years) in the GENEVA data using logistic regression under an additive genetic model, adjusting for gender and the top 3 principal components of ancestry (Manhattan and quantile-quantile plots are shown in Figures S4 and S5, respectively). Interestingly, we observed the same lung cancer risk SNP *rs12914385* also yielded a strong association with the dichotomized pack-year variable (*p=1.364×10^−6^*) and likewise was the top signal in the chromosome 15q25.1 locus (see Figure S6).

Based on our results, we explored the relationship of *rs12914385* with smoking pack-years and lung cancer within a mediation framework. Specifically, we were interested in exploring how much this SNP (exposure *A*) influenced lung cancer disease status (outcome *Y*) through the dichotomized smoking pack-year variable (mediator *M*). To be consistent with the GWAS analyses, we assumed an additive genetic model for the SNP and thus coded *A* as the number of copies of the minor allele a subject possessed. Our analyses also controlled for gender and the top three principal components of ancestry (covariates *C*). Within controls, the minor-allele frequency of *rs12914385* was 0.402. We performed inference using our likelihood approach as well as the original VW method and the IPW-VW approach. For the IPW-VW approach, we assumed an disease prevalence of 6% which mirrors results reported by the American Cancer Society^36^. We note that our proposed mediation analyses for estimating the direct and indirect effects of SNP on lung cancer requires the no-unmeasured-confounding assumptions (I)-(IV) previously stated. We note that we can’t rule out that exposure, mediator, and outcome might be influenced by some latent phenotype (e.g. another cancer subtype) that the study did not collect.

We note that VanderWeele and colleagues^17,37^ previously conducted similar analyses to untangle the relationship between SNPs in 15q25.1 locus, smoking, and lung cancer using case-control mediation tools. Our analysis above differs from these previous analyses in a few important ways. First, the previous analysis did not use the GENEVA study and instead analyzed a different collection of genetic studies of lung cancer. Second, the authors did not consider *rs12914385* as the SNP exposure and instead studied two nearby SNPs (*rs8034191, rs1051730*) in moderate linkage disequilibrium with *rs12914385*. Finally, they also considered different measures of smoking than pack-years and did not adjust for principal components of ancestry as we do in our analyses presented here.

## RESULTS

### Simulations with Continuous Mediator

Table 2 provides simulation results for estimating and conducting inference of *OR^TE^, OR^NDE^, OR^NIE^, OR^CDE^* using our likelihood approach, the original VW method, and the VW-IPW under the generating model that assumed a continuous mediator with no mediator-exposure interaction effect on disease risk. We noticed important differences among the methods with regards to inference of indirect effects (highlighted in Table 2). In particular, the simulation results revealed that our likelihood approach yielded smaller (but still well calibrated) standard errors of the indirect effects, relative to the VW approach, due in part to the former method incorporating cases within the mediation model that the latter method discards. Consequently, Table 2 shows our likelihood approach has greater power to detect the indirect effect relative to the VW method. We performed additional power simulations under a wider range of indirect-effect size estimates (generated by varying the value of *γ_M_* in (2c) while holding the remaining simulation parameters at the values shown in Table 1) and observed that the likelihood approach was substantially more powerful than the VW method for detecting an indirect effect across all values considered (Figure 2). For total and direct effects, the likelihood, VW, and VW-IPW methods all yielded similar findings.

**TABLE 2:** CONTINUOUS MEDIATOR WITH NO INTERACTION.

| | | | | | | | Power at $\alpha =$ | | |
| --- | --- | --- | --- | --- | --- | --- | --- | --- | --- |
|  |  | True Value | Mean Value | Mean SE | SD | 95% CI Coverage | 0.05 | 0.01 | 0.001 |
| **Likelihood** |  |  |  |  |  |  |  |  |  |
| | $\log(OR^{TE})$ | **0.145** | 0.144 | 0.083 | 0.081 | 0.961 | 0.408 | 0.198 | 0.055 |
| | $\log(OR^{NDE})$ | **0.100** | 0.099 | 0.085 | 0.083 | 0.958 | 0.199 | 0.072 | 0.014 |
| | $\log(OR^{NIE})$ | **0.045** | **0.045** | **0.019** | **0.019** | **0.956** | **0.691** | **0.362** | **0.058** |
| | $\log(OR^{CDE})$ | **0.100** | 0.099 | 0.085 | 0.083 | 0.958 | 0.199 | 0.072 | 0.014 |
| **VW** |  |  |  |  |  |  |  |  |  |
| | $\log(OR^{TE})$ | **0.145** | 0.144 | 0.084 | 0.081 | 0.964 | 0.397 | 0.182 | 0.046 |
| | $\log(OR^{NDE})$ | **0.100** | 0.099 | 0.085 | 0.083 | 0.958 | 0.199 | 0.072 | 0.014 |
| | $\log(OR^{NIE})$ | **0.045** | **0.045** | **0.022** | **0.021** | **0.940** | **0.504** | **0.131** | **0.009** |
| | $\log(OR^{CDE})$ | **0.100** | 0.099 | 0.085 | 0.083 | 0.958 | 0.199 | 0.072 | 0.014 |
| **VW-IPW** |  |  |  |  |  |  |  |  |  |
| | $\log(OR^{TE})$ | **0.145** | 0.144 | 0.084 | 0.081 | 0.964 | 0.397 | 0.185 | 0.047 |
| | $\log(OR^{NDE})$ | **0.100** | 0.099 | 0.085 | 0.083 | 0.958 | 0.199 | 0.072 | 0.014 |
| | $\log(OR^{NIE})$ | **0.045** | **0.045** | **0.022** | **0.021** | **0.948** | **0.521** | **0.143** | **0.008** |
| | $\log(OR^{CDE})$ | **0.100** | 0.099 | 0.085 | 0.083 | 0.958 | 0.199 | 0.072 | 0.014 |

**FIGURE 2:**
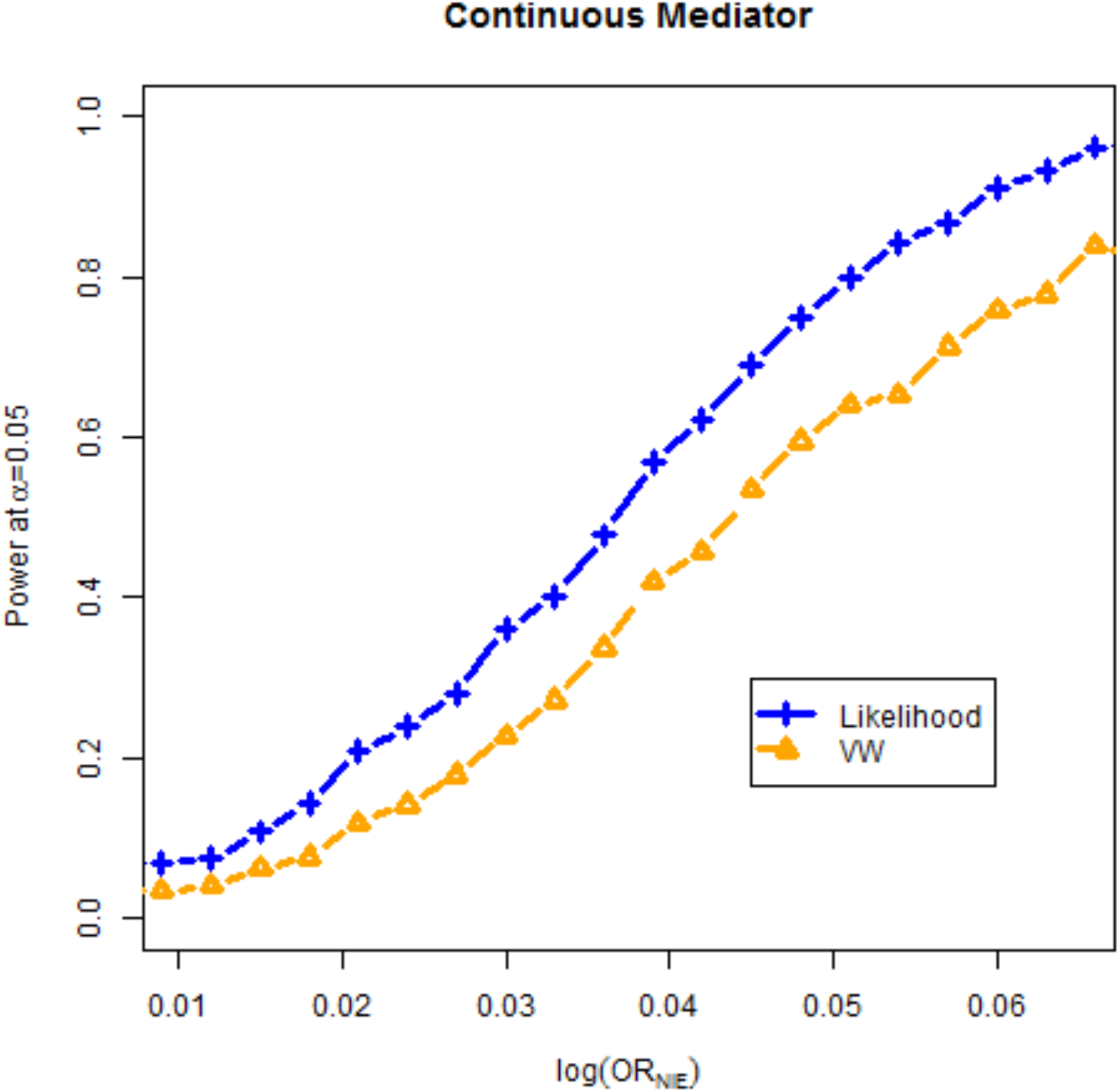
POWER TO DETECT INDIRECT EFFECT (CONTINUOUS MEDIATOR) Power of likelihood framework and original VW method to detect log (*OR^NIE^*) for 300 cases and 300 controls assuming continuous mediator and no exposure-mediator interaction effect. Simulation parameters shown in Table 1 for continuous mediator, with exception of *γ_M_* which is varied to produce range of values of log (*OR^NIE^*) shown in x-axis of plot.

Our likelihood approach improves efficiency of indirect effects by incorporating cases within the mediation model. This impact can be illuminated by studying the standard errors of the mediator-model parameter estimates (see Table S1), which are much smaller for our likelihood approach than for the VW approach. Additionally, by jointly modeling disease, mediator, and exposure data in a unified framework, we account for correlations among the disease-model and mediator-model parameters (see Table S2); the VW framework assumes no correlation among parameters from these two different models. By incorporating these non-zero correlations in our likelihood framework, our approach refines the standard errors of the indirect effects estimated using the delta method. Results using the VW-IPW approach (which incorporates cases but substantially downweights their contribution relative to controls) were quite similar to those using the VW approach but did yield slightly smaller standard errors for mediator-model parameter estimates relative to VW.

We also considered simulation models where we generated a mediator-exposure interaction and present the estimates and standard errors for *OR^TE^, OR^NDE^, OR^NIE^, OR^CDE^* in Table S3. For indirect effects, the VW and VW-IPW methods had higher coverage for the 95% CI than the nominal level (98.0% for both the VW and VW-IPW methods). Our likelihood approach, on the other hand, had appropriate coverage of the 95% confidence interval for the indirect effect. Consequently, the likelihood approach had increased power to detect the indirect effect relative to the VW and VW-IPW techniques. Inspection of the parameter estimates (Table S4) show that the likelihood approach provides more efficient estimates of the mediator-exposure interaction effect in the disease model, as well as those parameters related to the mediator model. Inspection of the mean covariance estimates for model parameters (Table S5) shows the likelihood approach estimates non-zero correlations between disease-model parameters (including the mediator-exposure interactions) and mediator-model parameters that the VW and IPW-VW methods presume are 0.

We next conducted additional simulations that assumed a disease prevalence of 20% to assess the impact of a common disease outcome on the methods considered. The results revealed our likelihood approach continued to have appropriate coverage for estimates of total, direct, and indirect effects and improved power relative to VW and VW-IPW methods (see Tables S6 and S7).

We finally performed type-I error simulations for testing the direct effect or indirect effect using the likelihood and VW approaches. For these simulations, we assumed a typical situation where one performs a mediation analysis where i) the exposure is known to be associated with disease outcome (i.e. *OR^TE^* ≠ 1) and ii) the exposure is known to be associated with mediator (i.e. *β*_A_, ≠ 0 in our model formulation). We first assumed no direct effect (*OR^NDE^* = 1) by using the same simulation setup shown in Table 1 but setting *γ_A_* = 0. Type I-error results across 10,000 simulations (shown in Table S8) revealed the likelihood approach and the VW methods have appropriate type-I error in this setting. We next conducted simulations assuming no indirect effect (*OR^NIE^* = 1) using the same simulation setup shown in Table 1 but setting *γ_M_* = 0. As shown in Table S8, we found that all three methods were conservative (with type-I error rates of 0.02-0.03 at alpha=0.05, depending on method), although the likelihood approach was better calibrated than VW techniques. The conservative nature of statistical tests of indirect effects have been identified by others^38,39^ and is due to formally testing a statistic under a composite null hypothesis (i.e. *H_O_*: *γ_M_β_A_* = 0) where the asymptotic distribution follows a mixture of normal product distributions (with unknown mixing probabilities) rather than a standard normal distribution.

### Simulations with Binary Mediator

Table 3 provides estimates and standard errors for *OR^TE^, OR^NDE^, OR^NIE^, OR^CDE^* for a binary mediator assuming no mediator-exposure interaction effect. For *OR^NIE^*, we observed that the likelihood approach yielded smaller standard errors than the VW and VW-IPW approaches and consequently had increased power to detect this indirect effect. We subsequently conducted additional power simulations under a broader range of indirect-effect size estimates (generated by varying values of *γ_M_* in (3c) while holding the remaining simulation parameters at the values shown in Table 1) and observed substantially increased power for the likelihood approach over the VW method across this range (Figure 3). The likelihood approach, VW, and VW-IPW approaches all produced similar results for total and direct effects. Inspection of parameter estimates (Table S9) and mean parameter covariance (Table S10) reveal similar trends to what we observed in the analysis of continuous mediators; the likelihood approach produces mediator-model parameter estimates with smaller standard errors than the VW and VW-IPW methods and the likelihood approach estimates non-zero covariances between disease-model parameters and mediator-model parameters.

**TABLE 3:** BINARY MEDIATOR WITH NO INTERACTION.

| | | | | | | | Power at $\alpha =$ | | |
| --- | --- | --- | --- | --- | --- | --- | --- | --- | --- |
|  |  | True Value | Mean Value | Mean SE | SD | 95% CI Coverage | 0.05 | 0.01 | 0.001 |
| **Likelihood** |  |  |  |  |  |  |  |  |  |
| | $\log(OR^{TE})$ | **0.145** | 0.142 | 0.083 | 0.083 | 0.953 | 0.391 | 0.193 | 0.061 |
| | $\log(OR^{NDE})$ | **0.100** | 0.097 | 0.085 | 0.085 | 0.949 | 0.200 | 0.080 | 0.021 |
| | $\log(OR^{NIE})$ | **0.045** | **0.045** | **0.020** | **0.021** | **0.940** | **0.649** | **0.319** | **0.034** |
| | $\log(OR^{CDE})$ | **0.100** | 0.097 | 0.085 | 0.085 | 0.949 | 0.200 | 0.080 | 0.021 |
| **VW** |  |  |  |  |  |  |  |  |  |
| | $\log(OR^{TE})$ | **0.145** | 0.142 | 0.084 | 0.083 | 0.961 | 0.373 | 0.184 | 0.055 |
| | $\log(OR^{NDE})$ | **0.100** | 0.097 | 0.085 | 0.085 | 0.949 | 0.200 | 0.080 | 0.021 |
| | $\log(OR^{NIE})$ | **0.045** | **0.045** | **0.023** | **0.024** | **0.920** | **0.497** | **0.134** | **0.008** |
| | $\log(OR^{CDE})$ | **0.100** | 0.097 | 0.085 | 0.085 | 0.949 | 0.200 | 0.080 | 0.021 |
| **VW-IPW** |  |  |  |  |  |  |  |  |  |
| | $\log(OR^{TE})$ | **0.145** | 0.142 | 0.084 | 0.083 | 0.961 | 0.374 | 0.185 | 0.055 |
| | $\log(OR^{NDE})$ | **0.100** | 0.097 | 0.085 | 0.085 | 0.949 | 0.200 | 0.080 | 0.021 |
| | $\log(OR^{NIE})$ | **0.045** | **0.045** | **0.022** | **0.023** | **0.923** | **0.519** | **0.153** | **0.008** |
| | $\log(OR^{CDE})$ | **0.100** | 0.097 | 0.085 | 0.085 | 0.949 | 0.200 | 0.080 | 0.021 |

**FIGURE 3:**
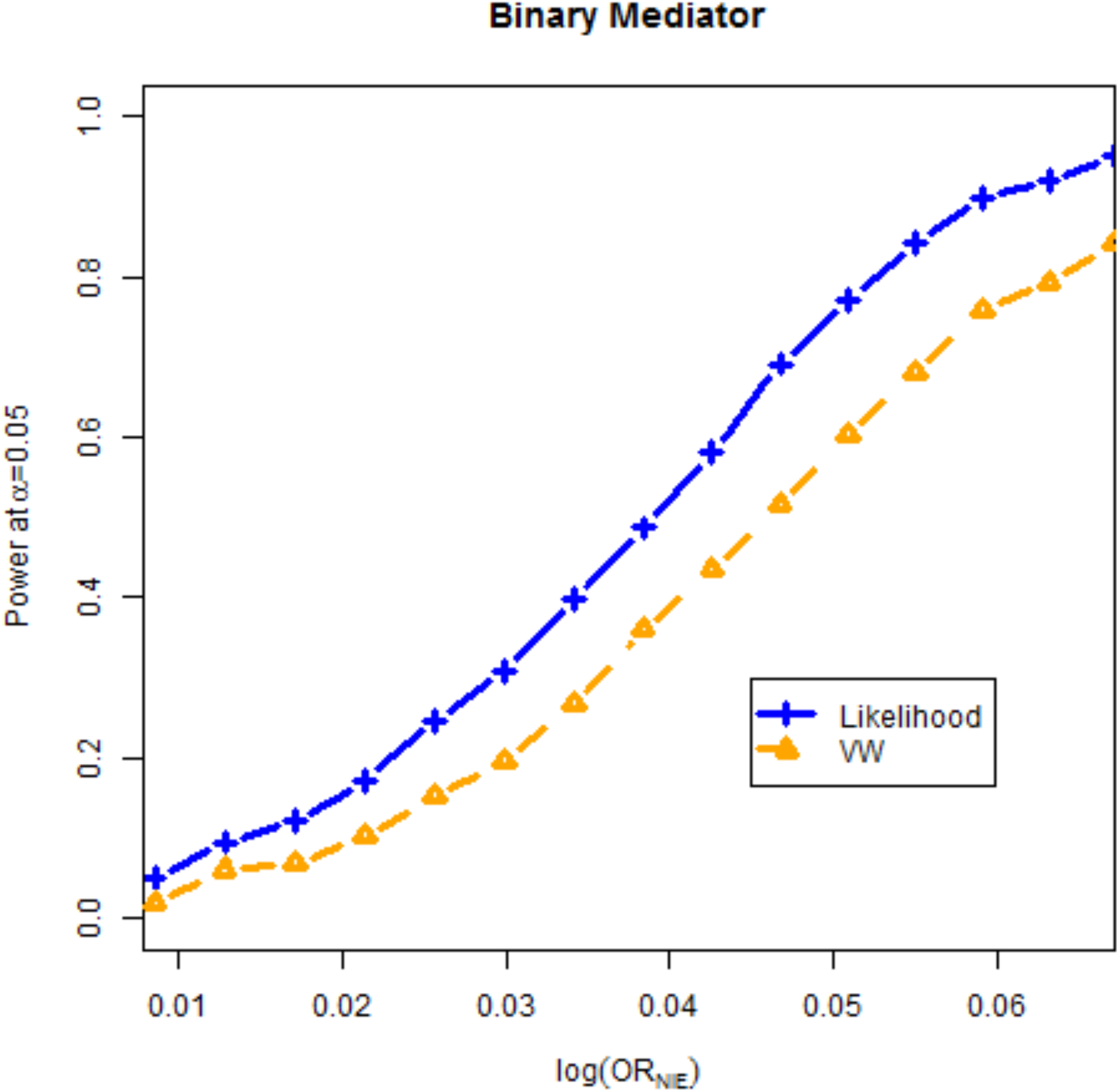
POWER TO DETECT INDIRECT EFFECT (BINARY MEDIATOR) Power of likelihood framework and original VW method to detect log (*OR^NIE^*) for 300 cases and 300 controls assuming binary mediator and no exposure-mediator interaction effect. Simulation parameters shown in Table 1 for binary mediator, with exception of *γ_M_* which is varied to produce range of values of log (*OR^NIE^*) shown in x-axis of plot.

We also considered simulation models with a binary mediator assuming a mediator-exposure interaction effect on disease risk and show the main results in Table S11. For the indirect effect, our likelihood approach had appropriate coverage of the 95% confidence interval while the VW and VW-IPW methods had higher coverage of this interval than the nominal level (98.6% for both methods). Consequently, the likelihood approach had markedly increased power to detect the indirect effect relative to the VW/VW-IPW methods. Inspection of model parameter estimates (Table S12) and mean parameter covariance (Table S13) revealed similar findings as previously observed for other simulation models. Results for direct and total effects were similar among all three methods when we modeled an interaction effect.

We conducted additional simulations assuming a more common disease outcome (prevalence of 20%) and observed similar trends to those that we observed under our original simulation models that assumed a disease prevalence of 3% (see Tables S14 and S15). We also performed type-I error simulations for direct and indirect effects (shown in Table S8) using the same design as for a continuous mediator. As with a continuous mediator, we observed that the likelihood and VW approaches had appropriate type-I error for testing the direct effect but were conservative when testing the indirect effect due to the evaluation of a composite null hypothesis.

### Mediation Analyses Ignoring Ascertainment

In evaluating the VW framework, we fit the original version of the method that fit the mediator model in equations (1b) and (1c) using only control data. As noted in the Introduction, naïvely fitting the mediation model on the combined case-control sample ignoring ascertainment can lead to bias as the disease prevalence decreases and the magnitudes of *γ_A_* and *γ_M_* increase. Since our simulation models in Table 1 assume a modest prevalence (0.03) and modest effects of *γ_A_* and *γ_M_*, we observed only slight bias in estimates of mediator-model parameters using the naïve method in our simulation results (Table S16) although we did observe naïve analysis led to weak coverage of indirect effect and had decreased power to detect this indirect effect compared to our likelihood approach (Table S17). To further interrogate the naïve method, we conducted additional simulations assuming a rarer prevalence (0.003) and larger effects for *γ_A_* (0.4) and *γ_M_* (0.6). In this setting, we observed more measurable bias in mediation-model parameters using the naïve approach while all other methods were unbiased (Table S18). We also observed that the naïve approach had even poorer coverage of the indirect effect than in our earlier simulations while all other methods had appropriate coverage (Table S19). The naïve approach also had substantially reduced power to detect the indirect effect compared to our likelihood approach. These results suggest one should avoid naïve combination of cases and controls (ignoring ascertainment) within the mediation model of traditional VW analyses.

### Application to Case-Control Genetic Study of Lung Cancer

Using the 2695 cases and 2779 controls from the GENEVA GWAS of lung cancer, we first conducted a mediation analysis that assessed whether the significant effect of SNP *rs12914385* on lung cancer risk was due in part to an indirect effect through a mediator of smoking pack-years, adjusting for covariate effects of gender and the top three principal components of ancestry. We initially applied the likelihood, VW, and IPW-VW approaches to the data explicitly assuming a potential interaction effect *γ_AM_* of exposure (SNP genotype) and mediator (smoking pack-years) on disease risk. However, we observed no significant interaction effect using the likelihood approach (*p=0.147*) or the VW/IPW-VW approaches (*p=0.130* for each). Thus, we removed the interaction parameter from the models and proceeded with reduced models that assumed no interaction effects.

Under the reduced models, we provide the odds-ratio estimates of indirect and direct effects of *rs12914385* on lung-cancer risk using our likelihood in Table 4 (see Table S20 for corresponding parameter estimates produced using our approach). Overall, our results demonstrate that the effect of *rs12914385* on lung cancer risk is predominantly a direct effect but there is evidence of a modest indirect effect through smoking pack-years. Specifically, the total effect of *rs12914385* on lung cancer risk (*OR^TE^* = 1.358; 95% CI: [1.255,1.470]) is nearly completely due to a direct effect (*OR^NDE^* =1.333; 95% CI: [1.226,1.449]) of the SNP on disease outcome rather than an indirect effect through the mediator of smoking pack-years (*OR^NIE^*=1.019; 95% CI: [1.000,1.039]). That being said, the test of the indirect effect was significant (*p=0.0494*) using our approach; additional analyses using larger sample sizes to explore the indirect effect further is warranted.

**TABLE 4:** MEDIATION ANALYSIS OF *rs12914385* AND LUNG CANCER.

|  |  | Estimate | 95% CI Coverage | P-value |
| --- | --- | --- | --- | --- |
| **Likelihood** |  |  |  |  |
| | $OR^{TE}$ | 1.358 | (1.255,1.470) | <0.0001 |
| | $OR^{NDE}$ | 1.333 | (1.226,1.449) | <0.0001 |
| | $OR^{NIE}$ | 1.019 | (1.000,1.039) | 0.0494 |
| | $OR^{CDE}$ | 1.333 | (1.226,1.449) | <0.0001 |
| **VW** |  |  |  |  |
| | $OR^{TE}$ | 1.347 | (1.232,1.474) | <0.0001 |
| | $OR^{NDE}$ | 1.333 | (1.226,1.449) | <0.0001 |
| | $OR^{NIE}$ | 1.011 | (0.978,1.044) | 0.5204 |
| | $OR^{CDE}$ | 1.333 | (1.226,1.449) | <0.0001 |
| **VW-IPW** |  |  |  |  |
| | $OR^{TE}$ | 1.359 | (1.244,1.484) | <0.0001 |
| | $OR^{NDE}$ | 1.333 | (1.226,1.450) | <0.0001 |
| | $OR^{NIE}$ | 1.019 | (0.991,1.048) | 0.1888 |
| | $OR^{CDE}$ | 1.333 | (1.226,1.450) | <0.0001 |

We next repeated the same mediation analysis of *rs129143851* using the traditional VW and IPW-VW approaches (Table 4, Table S20). While the estimates of total, direct, and indirect effects using the traditional VW approach were quite similar to those produced using our likelihood approach, we observed that the VW method’s confidence intervals were noticeably wider both for the total effect as well as the indirect effect compared to our likelihood approach. Furthermore, the VW approach found no borderline evidence of an indirect effect of *rs129143851* on lung-cancer risk through smoking pack-years (*p=0.5204*). Applying the IPW-VW approach to the dataset yielded similar point estimates of total, direct, and indirect effects as our likelihood approach. However, the 95% confidence intervals for the VW-IPW approach were wider than the analogous intervals for the likelihood approach for both the indirect effect and the total effect. Moreover, unlike the likelihood approach, the indirect effect estimate using VW-IPW was not significant (*p*=0.188).

## DISCUSSION

Our work develops a unified likelihood approach for case-control mediation analysis that improves inference of indirect effects relative to the popular counterfactual framework of VanderWeele by jointly modeling the disease model and mediator model together in a joint framework that further leverages important information on the mediator-exposure relationship within cases. Existing VW methods either ignore cases in the mediator model or use IPW procedures to incorporate them into the model. Unlike our approach, IPW requires knowledge of disease prevalence which may be difficult to ascertain. Even if prevalence information is correctly specified, the IPW-VW techniques fit the disease model and mediator model separately and so does not account for the correlation among parameters between the two models. Our likelihood framework utilizes this correlation, thereby resulting in more efficient estimates of parameters within the mediator model compared to the VW models. We also note our framework also leads to more efficient estimates of the mediator-exposure interaction variable in the disease model when modeled. Due to the increased efficiency of these parameter estimates (as well as accounting for parameter correlation between disease and mediator models), we also subsequently obtain more efficient estimates of indirect effects between an exposure and disease outcome compared to the VW framework. Our method further does not require knowledge of disease prevalence to incorporate cases into our framework.

We illustrated our method using genetic data from a GWAS study of lung cancer and found that the effect of the top risk SNP in the chromosome 15q25.1 locus on lung cancer within this dataset is predominantly a direct effect but there is some evidence for a modest indirect effect through smoking pack-years. While we applied our unified likelihood within a case-control genetics project for illustration purposes, the technique is widely applicable in other settings (e.g. environmental studies, psychological studies) that employ a case-control sampling design. We provide R code implementing the likelihood approach on our website (see Data Availability Statement).

Unlike indirect effects, we observed that our likelihood approach had similar performance for testing direct effects compared to VW methods. This result arises because, when there is no mediator-exposure interaction effect, the standard error of the direct effect (on log scale) depends only on the estimated variance of the exposure effect on disease (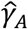 within disease-odds model in 1a). Since both the likelihood and VW methods estimate this quantity using the complete dataset, they yield similar standard errors for the estimated direct effect and therefore have similar power. If mediator-exposure interaction is modeled, the formula for calculating the standard errors of the direct effects becomes more complicated and depends on the estimated variance/covariances of several parameters (see Supplemental Materials) but, in most practical settings, these quantities are similar between the methods.

Power to detect the total effect that exposure has on disease depends on both the power to detect direct effect as well as the power to detect indirect effect. In our simulations in Table 1, we assumed the total effect was driven predominantly by direct effect rather than indirect effect. As the power of our likelihood to detect direct effects is the same as that of the VW approach, this results in our likelihood approach having a negligible power increase over the VW approach to detect the total effect in this particular situation. However, if we modify our simulations such that the total effect is driven predominantly by indirect effect rather than direct effect, then we do see a power increase of our likelihood approach to detect total effects compared to the VW approach (results not shown).

While the focus of this work was on traditional case-control mediation analysis for a continuous or binary exposure based on a counterfactual framework, we can potentially expand our likelihood framework to consider other current topics in this field. For example, there is substantial interest in jointly modeling multiple (correlated) mediators in case-control studies. VanderWeele and Vansteelandt^12^ expanded the standard VW framework to handle multiple mediators while Wang et al.^40^ developed a separate regression approach for assessing indirect effects that allows for one mediator to causally influence another mediator and further can correct for case-control ascertainment using a system of non-linear equations provided disease prevalence is known. We can expand our likelihood framework to handle multiple mediators by replacing the scalar mediator model in (1b) and (1c) with a multivariate mediator distribution for the controls under a rare disease approximation. Our approach would use cases in the mediator model (whereas the corresponding VW model^12^ does not) and does not require knowledge of disease prevalence (unlike Wang et al.).

We only considered the standard (unmatched) case-control design in this work, but note that we can potentially expand our likelihood approach to handle matched case-control datasets, such as those that arise in nested case-control studies. The matched design still allows estimates of direct and indirect effects conditional on the matching covariates^41,42^, where parameter estimates for the disease-odds model relating disease to mediator and exposure can be derived based on conditional logistic regression using the work of Kim et al^42^. We previously developed a retrospective likelihood for testing gene and gene-environment interactions in matched case-control designs^43^ and can implement a similar procedure based on that framework for this topic.

Our work focused on the simple situation in case-control mediation analysis where there is no missing information and the exposure is a scalar variable. However, practical situations can arise where a mediator may be censored; for example, a study may seek to investigate effects of genetic variants on breast cancer risk potentially mediated by (partially observed) age of menarche or age of menopause. We can explore a variation of our approach to handle censored mediators utilizing accelerated failure time models as described by Wang et al.^44,45^. We can also seek to expand our method for mediation analysis to handle a nominal multicategorical exposure variable^46^ (for example, a multi-SNP haplotype). This would require replacing the standard coding for *A* in equations (1a), (1b), and (1c) with a vector that represents dummy coding relating each categorical observation to a baseline category. Wang et al. ^45^ derived formulas for calculating the overall direct and indirect effects of the multicategorical exposure in this situation.

Both our likelihood approach and the VW framework model the probability of disease conditional on exposure and mediator even though, by design, case-control studies generate retrospectively ascertained mediator and exposure data. Such prospective analysis can be as efficient as a retrospective analysis but this result holds only under the assumption of a saturated non-parametric distribution for the exposure^20^. Carroll et al. ^47^ showed that prospective analysis of retrospectively-sampled data may be less efficient than a retrospective analysis when one restricts the exposure distribution in some fashion. For example, genetic studies often restrict the distribution of exposure (genotype or haplotype) by making the logical assumption that the underlying alleles are in Hardy-Weinberg Equilibrium (HWE). In these situations, many studies have shown^43,48–50^ that retrospective analysis of case-control genetic data can be more efficient than prospective analysis under the HWE assumption. Thus, in future work, we will investigate the use of a retrospective likelihood for case-control mediation analysis that may improve on prospective approaches by directly modeling the distribution of exposure conditional on outcome.

## ACKNOWLEDGEMENTS

This work was supported by NIH grants DE029698 and GM117946.

## DATA AVAILABILITY STATEMENT

Public R software implementing the likelihood approach can be found on the Epstein Software page (https://sites.google.com/view/epsteinlab/software). Simulation code is available on request from the authors. The GWAS lung-cancer dataset can be requested from the dbGaP site for the GENEVA study (https://www.ncbi.nlm.nih.gov/projects/gap/cgi-bin/study.cgi?study_id=phs000093.v2.p2)

## BIBLIOGRAPHY

1. MacKinnon D. Introduction to statistical mediation analysis. Routledge; 2012.

2. MacKinnon DP. Integrating mediators and moderators in research design. Research on social work practice. 2011;21(6): 675–681.

3. VanderWeele TJ. Mediation analysis: a practitioners guide. Annual review of public health. 2016;37:17–32.

4. Richiardi L, Bellocco R, Zugna D. Mediation analysis in epidemiology: methods, interpretation and bias. International journal of epidemiology. 2013;42(5): 1511–1519.

5. Baron RM, Kenny DA. The moderator-mediator variable distinction in social psychological research: Conceptual, strategic, and statistical considerations. Journal of personality and social psychology. 1986;51(6): 1173.

6. Judd CM, Kenny DA. Process analysis: Estimating mediation in treatment evaluations. Evaluation review. 1981;5(5): 602–619.

7. Sobel ME. Asymptotic confidence intervals for indirect effects in structural equation models. Sociological methodology. 1982;13:290–312.

8. Pearl J. Direct and indirect effects. Morgan Kaufmann Publishers Inc.; 2001:411–420.

9. Robins JM, Greenland S. Identifiability and exchangeability for direct and indirect effects. Epidemiology. 1992:143–155.

10. Valeri L, VanderWeele TJ. Mediation analysis allowing for exposure-mediator interactions and causal interpretation: theoretical assumptions and implementation with SAS and SPSS macros. Psychological methods. 2013;18(2): 137.

11. VanderWeele TJ, Vansteelandt S. Odds ratios for mediation analysis for a dichotomous outcome. American journal of epidemiology. 2010;172(12): 1339–1348.

12. VanderWeele T, Vansteelandt S. Mediation analysis with multiple mediators. Epidemiologic methods. 2014;2(1): 95–115.

13. Monsees GM, Tamimi RM, Kraft P. Genome - wide association scans for secondary traits using case - control samples. Genetic Epidemiology: The Official Publication of the International Genetic Epidemiology Society. 2009;33(8): 717–728.

14. Lin D, Zeng D. Proper analysis of secondary phenotype data in case - control association studies. Genetic Epidemiology: The Official Publication of the International Genetic Epidemiology Society. 2009;33(3): 256–265.

15. Wang J, Shete S. Estimation of odds ratios of genetic variants for the secondary phenotypes associated with primary diseases. Genetic epidemiology. 2011;35(3): 190–200.

16. Richardson DB, Rzehak P, Klenk J, Weiland SK. Analyses of case-control data for additional outcomes. Epidemiology. 2007;18(4):441–445.

17. VanderWeele TJ, Asomaning K, Tchetgen Tchetgen EJ, et al. Genetic variants on 15q25. 1, smoking, and lung cancer: an assessment of mediation and interaction. American journal of epidemiology. 2012;175(10): 1013–1020.

18. White H. A heteroskedasticity–consistent covariance matrix estimator and a direct test for heteroskedasticity. Econometrica: journal of the Econometric Society. 1980:817–838.

19. Huber PJ. The behavior of maximum likelihood estimates under nonstandard conditions. University of California Press; 1967:221–233.

20. Prentice RL, Pyke R. Logistic disease incidence models and case-control studies. Biometrika. 1979;66(3): 403–411.

21. Satten GA, Kupper LL. Inferences about exposure-disease associations using probability-of-exposure information. Journal of the American Statistical Association. 1993;88(421):200–208.

22. Satten GA, Carroll RJ. Conditional and unconditional categorical regression models with missing covariates. Biometrics. 2000;56(2): 384–388.

23. Siegel R, Ma J, Zou Z, Jemal A. Cancer statistics, 2014. CA: a cancer journal for clinicians. 2014;64(1):9–29.

24. Humans IWGotEoCRt, Organization WH, Cancer IAfRo. Tobacco smoke and involuntary smoking. Iarc; 2004.

25. Jonsson S, Thorsteinsdottir U, Gudbjartsson DF, et al. Familial risk of lung carcinoma in the Icelandic population. Jama. 2004;292(24): 2977–2983.

26. Mucci LA, Hjelmborg JB, Harris JR, et al. Familial risk and heritability of cancer among twins in Nordic countries. Jama. 2016;315(1): 68–76.

27. McKay JD, Hung RJ, Han Y, et al. Large-scale association analysis identifies new lung cancer susceptibility loci and heterogeneity in genetic susceptibility across histological subtypes. Nature genetics. 2017;49(7): 1126–1132.

28. Bossé Y, Amos CI. A decade of GWAS results in lung cancer. Cancer Epidemiology and Prevention Biomarkers. 2018;27(4): 363–379.

29. Hung RJ, McKay JD, Gaborieau V, et al. A susceptibility locus for lung cancer maps to nicotinic acetylcholine receptor subunit genes on 15q25. Nature. 2008;452(7187):633–637.

30. Freathy RM, Ring SM, Shields B, et al. A common genetic variant in the 15q24 nicotinic acetylcholine receptor gene cluster (CHRNA5-CHRNA3-CHRNB4) is associated with a reduced ability of women to quit smoking in pregnancy. Human molecular genetics. 2009;18(15): 2922–2927.

31. Munafò MR, Johnstone EC, Walther D, Uhl GR, Murphy MF, Aveyard P. CHRNA3 rs1051730 genotype and short-term smoking cessation. Nicotine & Tobacco Research. 2011;13(10):982–988.

32. Saccone NL, Culverhouse RC, Schwantes-An T-H, et al. Multiple independent loci at chromosome 15q25. 1 affect smoking quantity: a meta-analysis and comparison with lung cancer and COPD. PLoS Genet. 2010;6(8):e1001053.

33. Weiss RB, Baker TB, Cannon DS, et al. A candidate gene approach identifies the CHRNA5-A3-B4 region as a risk factor for age-dependent nicotine addiction. PLoS Genet. 2008;4(7): e1000125.

34. Prorok PC, Andriole GL, Bresalier RS, et al. Design of the prostate, lung, colorectal and ovarian (PLCO) cancer screening trial. Controlled clinical trials. 2000;21(6):273S–309S.

35. Landi MT, Consonni D, Rotunno M, et al. Environment And Genetics in Lung cancer Etiology (EAGLE) study: an integrative population–based case–control study of lung cancer. BMC public health. 2008;8(1): 1–10.

36. Society AC. Key Statistics for Lung Cancer. Accessed October 19, 2021, https://www.cancer.org/cancer/lung-cancer/about/key-statistics.html

37. VanderWeele TJ. A three-way decomposition of a total effect into direct, indirect, and interactive effects. Epidemiology (Cambridge, Mass) 2013;24(2): 224.

38. Barfield R, Shen J, Just AC, et al. Testing for the indirect effect under the null for genome - wide mediation analyses. Genetic epidemiology. 2017;41(8): 824–833.

39. Huang Y-T. Genome-wide analyses of sparse mediation effects under composite null hypotheses. The Annals of Applied Statistics. 2019;13(1): 60–84.

40. Wang J, Spitz MR, Amos CI, et al. Method for evaluating multiple mediators: mediating effects of smoking and COPD on the association between the CHRNA5-A3 variant and lung cancer risk. 2012;

41. VanderWeele TJ, Tchetgen Tchetgen EJ. Mediation analysis with matched case-control study designs. American journal of epidemiology. 2016;183(9):869–870.

42. Kim YM, Cologne JB, Jang E, et al. Causal mediation analysis in nested case - control studies using conditional logistic regression. Biometrical Journal. 2020;62(8): 1939–1959.

43. Kwee L, Epstein M, Manatunga A, Duncan R, Allen A, Satten G. Simple methods for assessing haplotype - environment interactions in case - only and case - control studies. Genetic Epidemiology: The Official Publication of the International Genetic Epidemiology Society. 2007;31(1): 75–90.

44. Wang J, Ning J, Shete S. Mediation analysis in a case - control study when the mediator is a censored variable. Statistics in medicine. 2019;38(7): 1213–1229.

45. Wang J, Ning J, Shete S. Mediation model with a categorical exposure and a censored mediator with application to a genetic study. PloS one. 2021;16(10):e0257628.

46. Hayes AF, Preacher KJ. Statistical mediation analysis with a multicategorical independent variable. British Journal of mathematical and statistical psychology. 2014;67(3): 451–470.

47. Carroll R, Wang S, Wang C. Prospective analysis of logistic case-control studies. Journal of the American Statistical Association. 1995;90(429): 157–169.

48. Epstein MP, Satten GA. Inference on haplotype effects in case-control studies using unphased genotype data. The American Journal of Human Genetics. 2003;73(6): 1316–1329.

49. Satten GA, Epstein MP. Comparison of prospective and retrospective methods for haplotype inference in case - control studies. Genetic Epidemiology: The Official Publication of the International Genetic Epidemiology Society. 2004;27(3): 192–201.

50. Chen J, Chatterjee N. Exploiting Hardy-Weinberg equilibrium for efficient screening of single SNP associations from case-control studies. Human heredity. 2007;63(3-4):196.

