## Supplementary files for "Efficient Estimation of Indirect Effects in Case-Control Studies Using a Unified Likelihood Framework"

### SUPPLEMENTARY METHODS

_Standard errors for direct and indirect effects for continuous mediator:_ We derive the standard errors of direct and indirect effects fit using our likelihood approach in (8) using the work of Valeri and VanderWeele[1]. We first assume we fit the likelihood in (8) assuming a continuous mediator, which yields parameter estimates $\hat{\gamma} = (\hat{\gamma}_0, \hat{\gamma}_A, \hat{\gamma}_M, \hat{\gamma}_{AM}, \hat{\gamma}_C')'$, $\hat{\beta} = \left(\hat{\beta}_0, \hat{\beta}_A, \hat{\beta}_C'\right)'$, and $\hat{\sigma}^2$ and estimated covariance matrix

$$\hat{\Sigma} = \begin{bmatrix} \Sigma_{\gamma,\gamma} & \Sigma_{\gamma,\beta} & \Sigma_{\gamma,\sigma^2} \\ \Sigma_{\beta,\gamma} & \Sigma_{\beta,\beta} & \Sigma_{\beta,\sigma^2} \\ \Sigma_{\sigma^2,\gamma} & \Sigma_{\sigma^2,\beta} & \Sigma_{\sigma^2,\sigma^2} \end{bmatrix}.$$

Based on these parameter estimates and estimated covariance matrix, we can derive the variance estimates for direct and indirect effects on the log scale outlined in equations (2a), (2b), and (2c) using the following formula based on the delta method

$$\Gamma\hat{\Sigma}\Gamma'(a - a^*)^2$$

Where $\Gamma$ denotes the vector of derivatives of $log\{OR^{NIE}\}$, $log\{OR^{NDE}\}$, or $log\{OR^{CDE}\}$ with respect to $(\gamma, \beta, \sigma^2)$. For $log\{OR^{CDE}\}$, $\Gamma = (0,1,0,m,0',0,0,0',0)$. For $log\{OR^{NDE}\}$, $\Gamma = \Big(0,1,\hat{\gamma}_{AM}\hat{\sigma}^2, \hat{\beta}_0 + \hat{\beta}_A a^* + \hat{\beta}_C c + \hat{\gamma}_M\hat{\sigma}^2 + \hat{\gamma}_{AM}\hat{\sigma}^2(a + a^*), 0', \hat{\gamma}_{AM}, \hat{\gamma}_{AM}a^*, \hat{\gamma}_{AM}c, \hat{\gamma}_M\hat{\gamma}_{AM} + 0.5\hat{\gamma}_{AM}^2(a + a^*)\Big)$. For $log\{OR^{NIE}\}$, $\Gamma = \left(0,0,\hat{\beta}_A, \hat{\beta}_A a, 0', 0, \hat{\gamma}_M + \hat{\gamma}_{AM}a, 0', 0\right)$. Finally, for $log\{OR^{TE}\} = log\{OR^{NDE}\} + log\{OR^{NIE}\}$, we have $\Gamma = \Big(0,1,\hat{\gamma}_{AM}\hat{\sigma}^2 + \hat{\beta}_A, \hat{\beta}_0 + \hat{\beta}_A(a + a^*) + \hat{\beta}_C c + \hat{\gamma}_M\hat{\sigma}^2 + \hat{\gamma}_{AM}\hat{\sigma}^2(a + a^*), 0', \hat{\gamma}_{AM}, \hat{\gamma}_M + \hat{\gamma}_{AM}(a + a^*), \hat{\gamma}_{AM}c, \hat{\gamma}_M\hat{\gamma}_{AM} + 0.5\hat{\gamma}_{AM}^2(a + a^*)\Big)$. We then calculate the standard error of each effect as the square root of its respective variance estimate.

_Standard errors for direct and indirect effects for binary mediator:_ We next derive the standard errors of direct and indirect effects fit using our likelihood approach in (8) assuming a binary mediator, which yields parameter estimates $\hat{\gamma} = (\hat{\gamma}_0, \hat{\gamma}_A, \hat{\gamma}_M, \hat{\gamma}_{AM}, \hat{\gamma}_C')'$ and $\hat{\beta} = \left(\hat{\beta}_0, \hat{\beta}_A, \hat{\beta}_C'\right)'$ and estimated covariance matrix

$$\hat{\Sigma} = \begin{bmatrix} \Sigma_{\gamma,\gamma} & \Sigma_{\gamma,\beta} \\ \Sigma_{\beta,\gamma} & \Sigma_{\beta,\beta} \end{bmatrix}.$$

Based on these parameter estimates and estimated covariance matrix, we can derive the variance estimate for direct and indirect effects on the log scale outlined in equations (3a), (3b), and (3c) using the following formula based on the delta method

$$\Gamma\hat{\Sigma}\Gamma'$$

Where $\Gamma$ denotes the vector of derivatives of $log\{OR^{NIE}\}$, $log\{OR^{NDE}\}$, or $log\{OR^{CDE}\}$ with respect to $(\gamma, \beta)$. For $log\{OR^{CDE}\}$, $\Gamma = (0, (a - a^*), 0, m(a - a^*), 0', 0,0, 0')$. For $log\{OR^{NDE}\}$, we first define $A = \frac{exp(\hat{\gamma}_M + \hat{\gamma}_{AM}a + \hat{\beta}_0 + \hat{\beta}_A a^* + \hat{\beta}_C c)}{1 + exp(\hat{\gamma}_M + \hat{\gamma}_{AM}a + \hat{\beta}_0 + \hat{\beta}_A a^* + \hat{\beta}_C c)}$ and $B = \frac{exp(\hat{\gamma}_M + \hat{\gamma}_{AM}a^* + \hat{\beta}_0 + \hat{\beta}_A a^* + \hat{\beta}_C c)}{1 + exp(\hat{\gamma}_M + \hat{\gamma}_{AM}a^* + \hat{\beta}_0 + \hat{\beta}_A a^* + \hat{\beta}_C c)}$ and then specify $\Gamma = \left(0, (a - a^*), A - B, aA - a^*B, 0', A - B, a^*(A - B), c'(A - B)\right)$. For $log\{OR^{NIE}\}$, we first define $A = \frac{exp(\hat{\gamma}_M + \hat{\gamma}_{AM}a + \hat{\beta}_0 + \hat{\beta}_A a + \hat{\beta}_C c)}{1 + exp(\hat{\gamma}_M + \hat{\gamma}_{AM}a + \hat{\beta}_0 + \hat{\beta}_A a + \hat{\beta}_C c)}$, $B = \frac{exp(\hat{\gamma}_M + \hat{\gamma}_{AM}a + \hat{\beta}_0 + \hat{\beta}_A a^* + \hat{\beta}_C c)}{1 + exp(\hat{\gamma}_M + \hat{\gamma}_{AM}a + \hat{\beta}_0 + \hat{\beta}_A a^* + \hat{\beta}_C c)}$, $K = \frac{exp(\hat{\beta}_0 + \hat{\beta}_A a + \hat{\beta}_C c)}{1 + exp(\hat{\beta}_0 + \hat{\beta}_A a + \hat{\beta}_C c)}$, and $D = \frac{exp(\hat{\beta}_0 + \hat{\beta}_A a^* + \hat{\beta}_C c)}{1 + exp(\hat{\beta}_0 + \hat{\beta}_A a^* + \hat{\beta}_C c)}$ and then specify $\Gamma = \Big(0,0, A - B, a(A - B), 0', (D + A) - (K + B), a^*(D - B) + a(A - K), c'\big((D + A) - (K + B)\big)\Big)$. Finally, for $log\{OR^{TE}\} = log\{OR^{NDE}\} + log\{OR^{NIE}\}$, we derive $\Gamma$ as the sum of the respective elements from those listed for $log\{OR^{NDE}\}$ and $log\{OR^{NIE}\}$. We then calculate the standard error of each effect as the square root of its respective variance estimate.

*Derivation of $\theta_j(a, c)$ for continuous mediator:* Assume our mediator follows a normal distribution $M_j \sim Normal(\beta_0 + \beta_A a + \beta_C c, \sigma^2)$. Based on equation (7), we can write $\theta_j(a, c)$ as

$$\theta_j(a,c) = exp(\gamma_0 + \gamma_A a + \gamma_C c) \int exp\big((\gamma_M + \gamma_{AM} a)m^*\big) \; * f_M(m^*)\, dm^* \quad \text{(A1)}$$

where $f_M(m^*) = \frac{1}{\sqrt{2\pi\sigma^2}} exp\left[-\frac{(m^* - \beta_0 - \beta_A a - \beta_C c)^2}{2\sigma^2}\right]$ is the relevant normal probability density function.

We can obtain a closed-form solution to the integrand in (A1) by leveraging knowledge of moment-generating functions for normal random variables. For a $X \sim Normal(\mu, \tau^2)$ variable, the moment-generating function for $X$ is $E[exp(zX)] = exp(\mu z + 0.5\tau^2 z^2)$. Assuming $M_j \sim Normal(\beta_0 + \beta_A a + \beta_C c, \sigma^2)$ and noting that the integrand in (A1) can be represented as $E\left[exp\left((\gamma_M + \gamma_{AM} a)M_j\right)\right]$, we can rewrite the integrand in (A1) as

$$\int exp((\gamma_M + \gamma_{AM} a)m^*) \; * f_M(m^*)\, dm^*$$

$$= exp\left((\beta_0 + \beta_A a + \beta_C c)(\gamma_M + \gamma_{AM} a) + 0.5\sigma^2(\gamma_M + \gamma_{AM} a)^2\right)$$

Substituting this closed-form solution for the integrand back into (A1), we obtain

$$\theta_j(a,c) = exp(\gamma_0 + \gamma_A a + (\gamma_M + \gamma_{AM} a)(\beta_0 + \beta_A a + \beta_C c) + 0.5(\gamma_M + \gamma_{AM} a)^2\sigma^2 + \gamma_C c)$$

### FIGURE S1: MANHATTAN PLOT FOR GENEVA GWAS OF LUNG CANCER

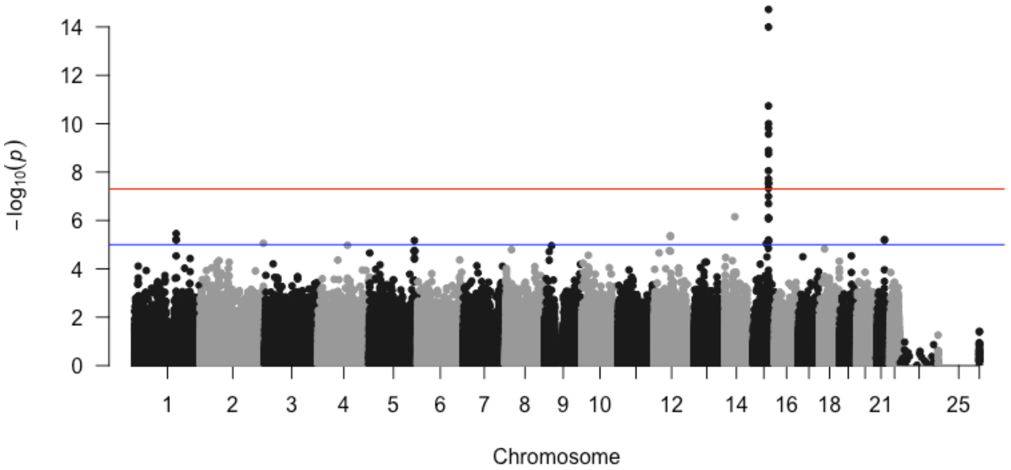

### FIGURE S2: QUANTILE-QUANTILE PLOT FOR GENEVA GWAS OF LUNG CANCER

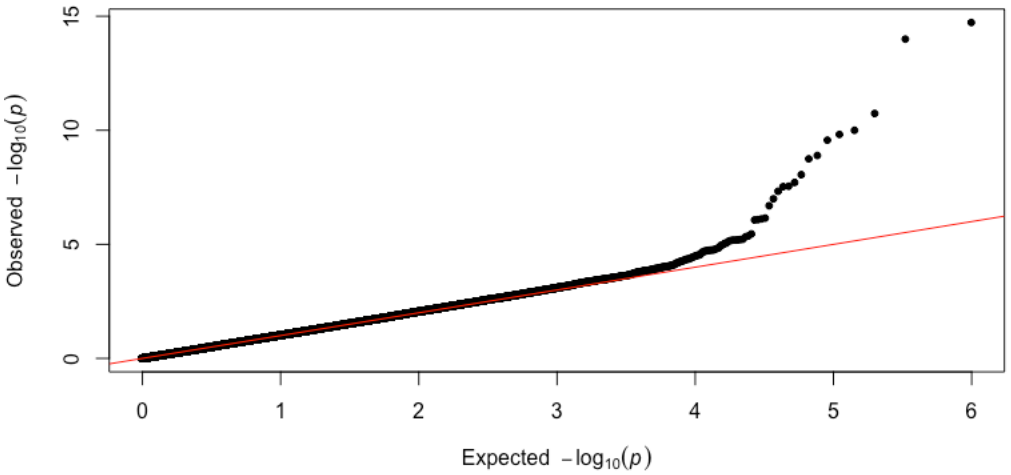

### FIGURE S3: ASSOCIATION OF CHROMOSOME 15q25.1 SNPS WITH LUNG CANCER

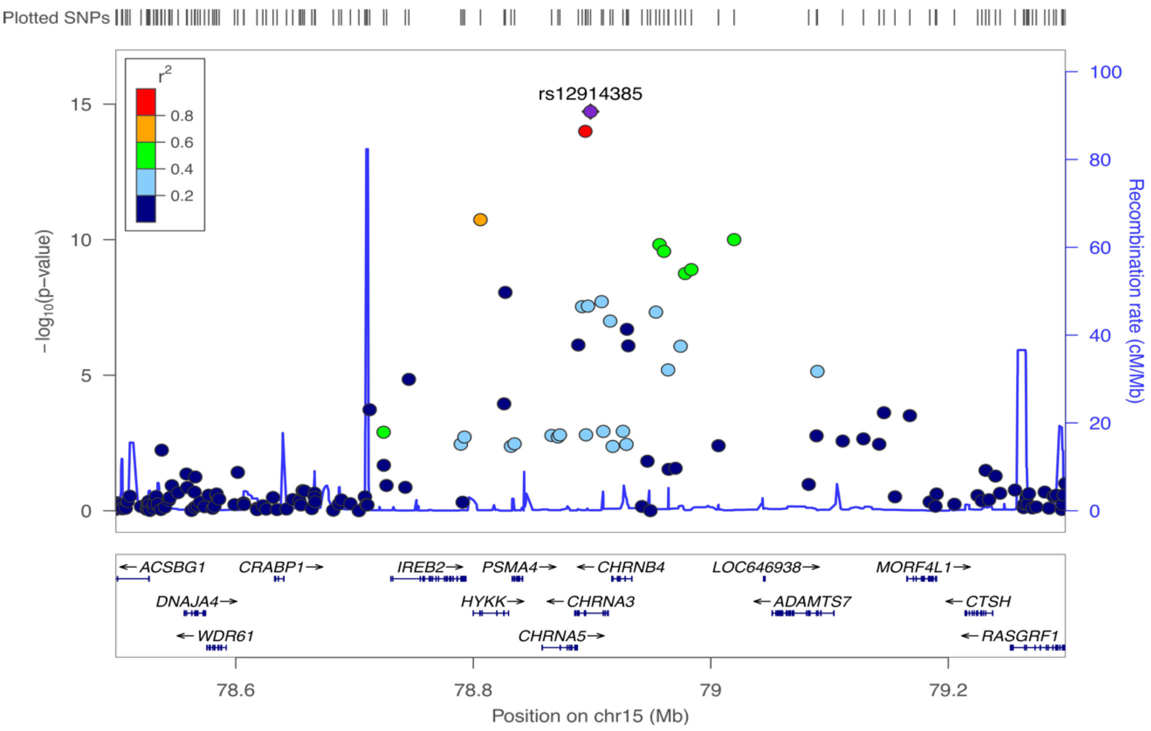

### FIGURE S4: MANHATTAN PLOT FOR GENEVA GWAS OF SMOKING PACK-YEARS (DICHOTOMOUS)

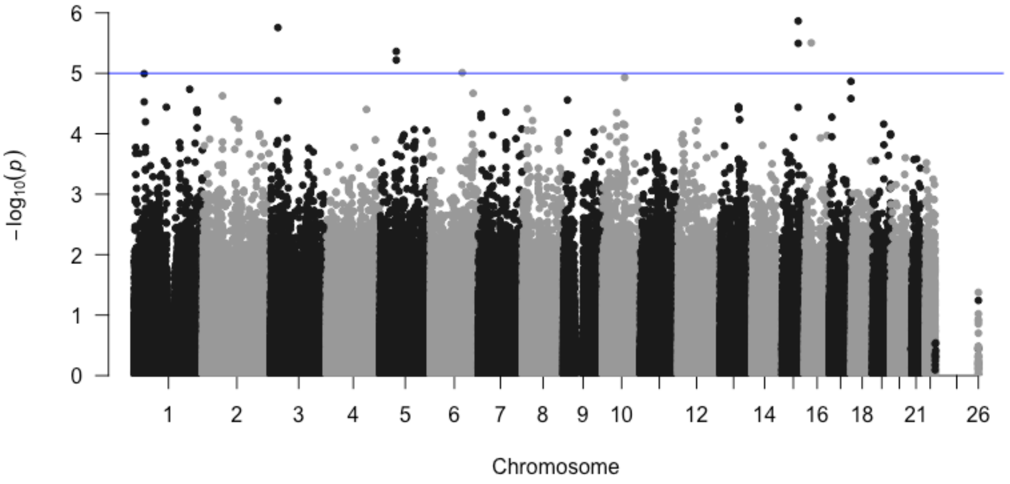

**FIGURE S5: QUANTILE-QUANTILE PLOT FOR GENEVA GWAS OF SMOKING PACK-YEARS (DICHOTOMOUS)**

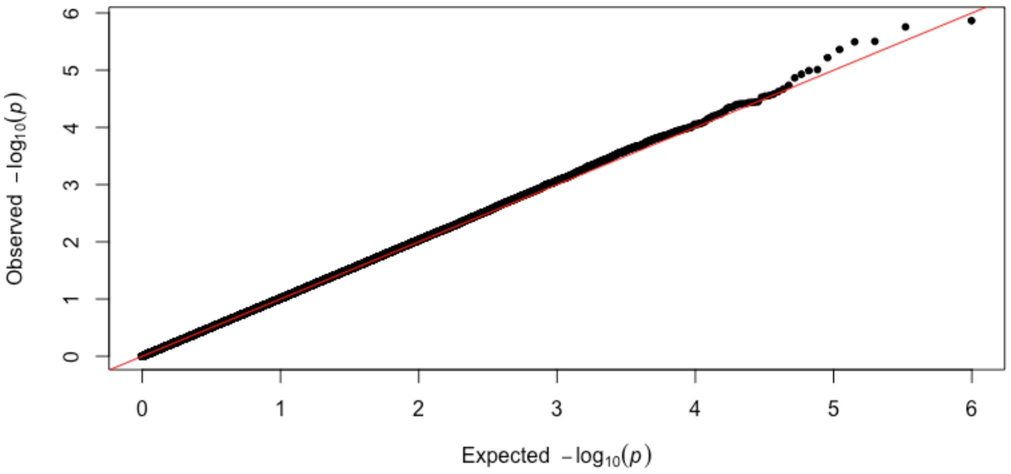

**FIGURE S6: ASSOCIATION OF CHROMOSOME 15q25.1 SNPS WITH SMOKING PACK-YEARS (DICHOTOMOUS)**

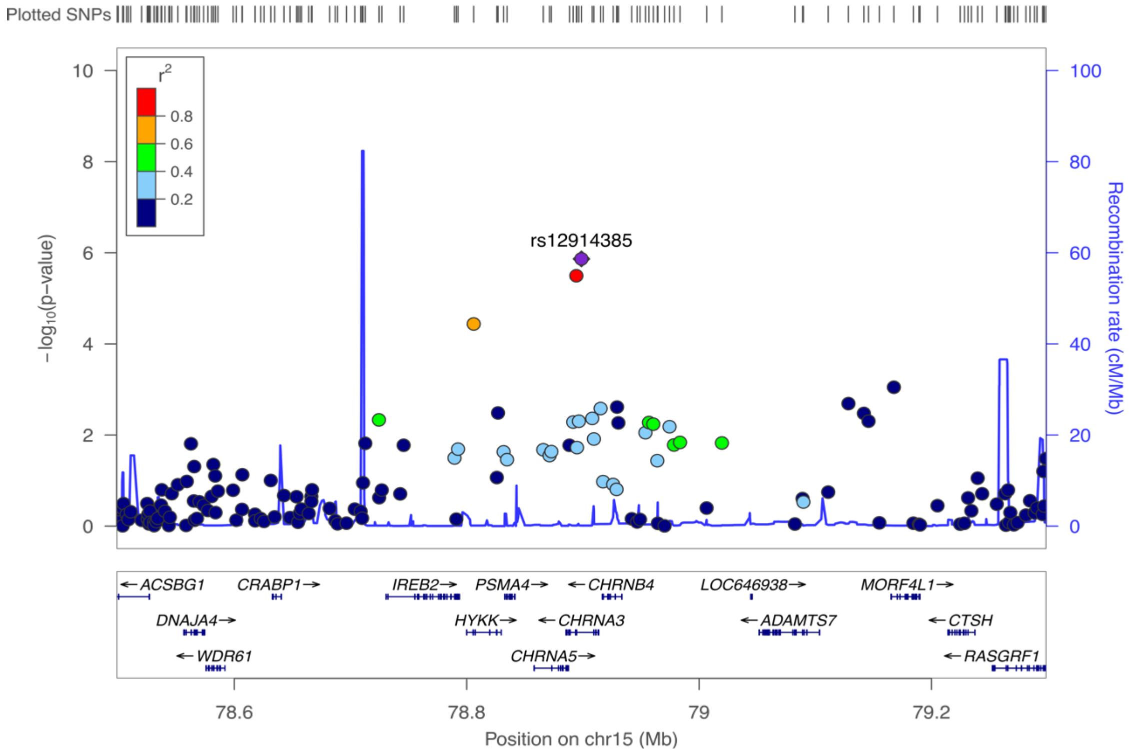

### TABLE S1: CONTINUOUS MEDIATOR WITH NO INTERACTION

| | True Value | Mean Value | Mean SE | SD |
|---|---|---|---|---|
| **Likelihood** | | | | |
| $\gamma_A$ | **0.100** | 0.099 | 0.085 | 0.083 |
| $\gamma_M$ | **0.300** | 0.304 | 0.118 | 0.114 |
| $\gamma_C$ | **0.050** | 0.052 | 0.166 | 0.167 |
| $\beta_0$ | **0.100** | 0.095 | 0.050 | 0.049 |
| $\beta_A$ | **0.150** | 0.149 | 0.029 | 0.028 |
| $\beta_C$ | **0.050** | 0.048 | 0.058 | 0.057 |
| $\sigma^2$ | **0.500** | 0.498 | 0.029 | 0.028 |
| **VW** | | | | |
| $\gamma_A$ | **0.100** | 0.099 | 0.085 | 0.083 |
| $\gamma_M$ | **0.300** | 0.304 | 0.118 | 0.114 |
| $\gamma_C$ | **0.050** | 0.052 | 0.166 | 0.167 |
| $\beta_0$ | **0.100** | 0.094 | 0.058 | 0.056 |
| $\beta_A$ | **0.150** | 0.149 | 0.041 | 0.040 |
| $\beta_C$ | **0.050** | 0.050 | 0.082 | 0.080 |
| $\sigma^2$ | **0.500** | 0.496 | 0.041 | 0.042 |
| **VW-IPW** | | | | |
| $\gamma_A$ | **0.100** | 0.099 | 0.085 | 0.083 |
| $\gamma_M$ | **0.300** | 0.304 | 0.118 | 0.114 |
| $\gamma_C$ | **0.050** | 0.052 | 0.166 | 0.167 |
| $\beta_0$ | **0.100** | 0.099 | 0.056 | 0.054 |
| $\beta_A$ | **0.150** | 0.150 | 0.040 | 0.039 |
| $\beta_C$ | **0.050** | 0.051 | 0.080 | 0.078 |
| $\sigma^2$ | **0.500** | 0.497 | 0.029 | 0.041 |

### TABLE S2: COVARIANCE ESTIMATES FOR CONTINUOUS MEDIATOR (NO INTERACTION)

| | | | | | | | |
|---|---|---|---|---|---|---|---|
| **Likelihood** | $\gamma_A$ | $\gamma_M$ | $\gamma_C$ | $\beta_0$ | $\beta_A$ | $\beta_C$ | $\sigma^2$ |
| $\gamma_A$ | 0.0072 | -0.002 | 0.0001 | 0.0005 | -0.0002 | 0 | 0.0001 |
| $\gamma_M$ | -0.002 | 0.0139 | -0.0006 | -0.0033 | -0.0002 | -0.0001 | -0.0005 |
| $\gamma_C$ | 0.0001 | -0.0006 | 0.0274 | 0.0007 | 0 | -0.001 | 0 |
| $\beta_0$ | 0.0005 | -0.0033 | 0.0007 | 0.0025 | 0 | -0.0017 | 0 |
| $\beta_A$ | -0.0002 | -0.0002 | 0 | 0 | 0.0008 | 0 | 0 |
| $\beta_C$ | 0 | -0.0001 | -0.001 | -0.0017 | 0 | 0.0033 | 0 |
| $\sigma^2$ | 0.0001 | -0.0005 | 0 | 0 | 0 | 0 | 0.0008 |
| **VW** | | | | | | | |
| $\gamma_A$ | 0.0072 | -0.002 | 0.0001 | 0 | 0 | 0 | 0 |
| $\gamma_M$ | -0.002 | 0.0139 | -0.0006 | 0 | 0 | 0 | 0 |
| $\gamma_C$ | 0.0001 | -0.0006 | 0.0274 | 0 | 0 | 0 | 0 |
| $\beta_0$ | 0 | 0 | 0 | 0.0033 | 0 | -0.0033 | 0 |
| $\beta_A$ | 0 | 0 | 0 | 0 | 0.0017 | 0 | 0 |
| $\beta_C$ | 0 | 0 | 0 | -0.0033 | 0 | 0.0067 | 0 |
| $\sigma^2$ | 0 | 0 | 0 | 0 | 0 | 0 | 0.0017 |
| **VW-IPW** | | | | | | | |
| $\gamma_A$ | 0.0072 | -0.002 | 0.0001 | 0 | 0 | 0 | 0 |
| $\gamma_M$ | -0.002 | 0.0139 | -0.0006 | 0 | 0 | 0 | 0 |
| $\gamma_C$ | 0.0001 | -0.0006 | 0.0274 | 0 | 0 | 0 | 0 |
| $\beta_0$ | 0 | 0 | 0 | 0.0032 | 0 | -0.0032 | 0 |
| $\beta_A$ | 0 | 0 | 0 | 0 | 0.0016 | 0 | 0 |
| $\beta_C$ | 0 | 0 | 0 | -0.0032 | 0 | 0.0064 | 0 |
| $\sigma^2$ | 0 | 0 | 0 | 0 | 0 | 0 | 0.0008 |

### TABLE S3: CONTINUOUS MEDIATOR WITH INTERACTION

| | True Value | Mean Value | Mean SE | Emp. SD | 95% CI Coverage | Power at $\alpha =$ 0.05 | 0.01 | 0.001 |
|---|---|---|---|---|---|---|---|---|
| **Likelihood** | | | | | | | | |
| $\log(OR^{TE})$ | **0.145** | 0.145 | 0.084 | 0.084 | 0.956 | 0.407 | 0.215 | 0.055 |
| $\log(OR^{NDE})$ | **0.100** | 0.103 | 0.085 | 0.086 | 0.951 | 0.240 | 0.086 | 0.018 |
| $\log(OR^{NIE})$ | **0.045** | 0.043 | 0.022 | 0.022 | 0.960 | 0.565 | 0.281 | 0.048 |
| $\log(OR^{CDE})$ | **0.077** | 0.079 | 0.087 | 0.090 | 0.940 | 0.143 | 0.057 | 0.006 |
| **VW** | | | | | | | | |
| $\log(OR^{TE})$ | **0.145** | 0.146 | 0.088 | 0.086 | 0.962 | 0.372 | 0.178 | 0.034 |
| $\log(OR^{NDE})$ | **0.100** | 0.104 | 0.086 | 0.087 | 0.952 | 0.238 | 0.087 | 0.020 |
| $\log(OR^{NIE})$ | **0.045** | 0.043 | 0.028 | 0.023 | 0.980 | 0.235 | 0.013 | 0.000 |
| $\log(OR^{CDE})$ | **0.077** | 0.079 | 0.088 | 0.091 | 0.941 | 0.152 | 0.054 | 0.005 |
| **VW-IPW** | | | | | | | | |
| $\log(OR^{TE})$ | **0.145** | 0.148 | 0.088 | 0.086 | 0.963 | 0.376 | 0.183 | 0.034 |
| $\log(OR^{NDE})$ | **0.100** | 0.104 | 0.086 | 0.087 | 0.952 | 0.239 | 0.088 | 0.020 |
| $\log(OR^{NIE})$ | **0.045** | 0.043 | 0.028 | 0.023 | 0.980 | 0.254 | 0.016 | 0.000 |
| $\log(OR^{CDE})$ | **0.077** | 0.079 | 0.088 | 0.091 | 0.941 | 0.152 | 0.054 | 0.005 |

### TABLE S4: CONTINUOUS MEDIATOR WITH INTERACTION

| | True Value | Mean Value | Mean SE | SD |
|---|---|---|---|---|
| **Likelihood** | | | | |
| $\gamma_A$ | **0.077** | 0.079 | 0.087 | 0.090 |
| $\gamma_M$ | **0.200** | 0.203 | 0.117 | 0.118 |
| $\gamma_{AM}$ | **0.100** | 0.099 | 0.109 | 0.112 |
| $\gamma_C$ | **0.050** | 0.054 | 0.165 | 0.168 |
| $\beta_0$ | **0.100** | 0.099 | 0.050 | 0.049 |
| $\beta_A$ | **0.150** | 0.148 | 0.040 | 0.041 |
| $\beta_C$ | **0.050** | 0.047 | 0.058 | 0.059 |
| $\sigma^2$ | **0.500** | 0.497 | 0.029 | 0.028 |
| **VW** | | | | |
| $\gamma_A$ | **0.077** | 0.079 | 0.088 | 0.091 |
| $\gamma_M$ | **0.200** | 0.204 | 0.118 | 0.119 |
| $\gamma_{AM}$ | **0.100** | 0.102 | 0.113 | 0.116 |
| $\gamma_C$ | **0.050** | 0.053 | 0.165 | 0.168 |
| $\beta_0$ | **0.100** | 0.099 | 0.058 | 0.057 |
| $\beta_A$ | **0.150** | 0.148 | 0.041 | 0.042 |
| $\beta_C$ | **0.050** | 0.047 | 0.082 | 0.083 |
| $\sigma^2$ | **0.500** | 0.497 | 0.041 | 0.039 |
| **VW-IPW** | | | | |
| $\gamma_A$ | **0.077** | 0.079 | 0.088 | 0.091 |
| $\gamma_M$ | **0.200** | 0.204 | 0.118 | 0.119 |
| $\gamma_{AM}$ | **0.100** | 0.102 | 0.113 | 0.116 |
| $\gamma_C$ | **0.050** | 0.053 | 0.165 | 0.168 |
| $\beta_0$ | **0.100** | 0.102 | 0.057 | 0.055 |
| $\beta_A$ | **0.150** | 0.150 | 0.040 | 0.041 |
| $\beta_C$ | **0.050** | 0.047 | 0.080 | 0.080 |
| $\sigma^2$ | **0.500** | 0.498 | 0.029 | 0.038 |

### TABLE S5: COVARIANCE ESTIMATES FOR CONTINUOUS MEDIATOR (WITH INTERACTION)

| **Likelihood** | | | | | | | | |
|---|---|---|---|---|---|---|---|---|
| | $\gamma_A$ | $\gamma_M$ | $\gamma_{AM}$ | $\gamma_C$ | $\beta_0$ | $\beta_A$ | $\beta_C$ | $\sigma^2$ |
| $\gamma_A$ | 0.0076 | -0.0022 | -0.0022 | 0.0001 | 0.0005 | 0.0004 | 0 | 0.0001 |
| $\gamma_M$ | -0.0022 | 0.0138 | -0.0007 | -0.0006 | -0.0033 | 0 | -0.0001 | -0.0003 |
| $\gamma_{AM}$ | -0.0022 | -0.0007 | 0.012 | 0 | 0 | -0.0031 | 0 | -0.0001 |
| $\gamma_C$ | 0.0001 | -0.0006 | 0 | 0.0273 | 0.0005 | 0 | -0.0007 | 0 |
| $\beta_0$ | 0.0005 | -0.0033 | 0 | 0.0005 | 0.0025 | 0 | -0.0017 | 0 |
| $\beta_A$ | 4.00E-04 | 0 | -0.0031 | 0 | 0 | 0.0016 | 0 | 0 |
| $\beta_C$ | 0 | -0.0001 | 0 | -0.0007 | -0.0017 | 0 | 0.0033 | 0 |
| $\sigma^2$ | 1.00E-04 | -0.0003 | -0.0001 | 0 | 0 | 0 | 0 | 0.0008 |
| **VW** | | | | | | | | |
| $\gamma_A$ | 0.0077 | -0.0022 | -0.0022 | 0.0001 | 0 | 0 | 0 | 0 |
| $\gamma_M$ | -0.0022 | 0.0139 | -0.0006 | -0.0006 | 0 | 0 | 0 | 0 |
| $\gamma_{AM}$ | -0.0022 | -0.0006 | 0.0127 | 0 | 0 | 0 | 0 | 0 |
| $\gamma_C$ | 0.0001 | -0.0006 | 0 | 0.0274 | 0 | 0 | 0 | 0 |
| $\beta_0$ | 0 | 0 | 0 | 0 | 0.0033 | 0 | -0.0033 | 0 |
| $\beta_A$ | 0 | 0 | 0 | 0 | 0 | 0.0017 | 0 | 0 |
| $\beta_C$ | 0 | 0 | 0 | 0 | -0.0033 | 0 | 0.0067 | 0 |
| $\sigma^2$ | 0 | 0 | 0 | 0 | 0 | 0 | 0 | 0.0017 |
| **VW-IPW** | | | | | | | | |
| $\gamma_A$ | 0.0077 | -0.0022 | -0.0022 | -0.0001 | 0 | 0 | 0 | 0 |
| $\gamma_M$ | -0.0022 | 0.0139 | -0.0006 | -0.0006 | 0 | 0 | 0 | 0 |
| $\gamma_{AM}$ | -0.0022 | -0.0006 | 0.0127 | 0 | 0 | 0 | 0 | 0 |
| $\gamma_C$ | 0.0001 | -0.0006 | 0 | 0.0274 | 0 | 0 | 0 | 0 |
| $\beta_0$ | 0 | 0 | 0 | 0 | 0.0032 | 0 | -0.0032 | 0 |
| $\beta_A$ | 0 | 0 | 0 | 0 | 0 | 0.0016 | 0 | 0 |
| $\beta_C$ | 0 | 0 | 0 | 0 | -0.0032 | 0 | 0.0064 | 0 |
| $\sigma^2$ | 0 | 0 | 0 | 0 | 0 | 0 | 0 | 0.0008 |

### TABLE S6: CONTINUOUS MEDIATOR, DISEASE PREVALENCE OF 0.20

| | True Value | Mean Value | Mean SE | Emp. SD | 95% CI Coverage | Power at $\alpha =$ 0.05 | 0.01 | 0.001 |
|---|---|---|---|---|---|---|---|---|
| **Likelihood** | | | | | | | | |
| $\log(OR^{TE})$ | **0.145** | 0.151 | 0.083 | 0.083 | 0.958 | 0.449 | 0.219 | 0.068 |
| $\log(OR^{NDE})$ | **0.100** | 0.107 | 0.085 | 0.085 | 0.955 | 0.237 | 0.082 | 0.015 |
| $\log(OR^{NIE})$ | **0.045** | 0.044 | 0.019 | 0.019 | 0.941 | 0.691 | 0.356 | 0.057 |
| $\log(OR^{CDE})$ | **0.100** | 0.107 | 0.085 | 0.085 | 0.955 | 0.237 | 0.082 | 0.015 |
| **VW** | | | | | | | | |
| $\log(OR^{TE})$ | **0.145** | 0.151 | 0.084 | 0.083 | 0.957 | 0.432 | 0.210 | 0.058 |
| $\log(OR^{NDE})$ | **0.100** | 0.107 | 0.085 | 0.085 | 0.955 | 0.237 | 0.082 | 0.015 |
| $\log(OR^{NIE})$ | **0.045** | 0.044 | 0.022 | 0.021 | 0.931 | 0.504 | 0.123 | 0.005 |
| $\log(OR^{CDE})$ | **0.100** | 0.107 | 0.085 | 0.085 | 0.955 | 0.237 | 0.082 | 0.015 |
| **VW-IPW** | | | | | | | | |
| $\log(OR^{TE})$ | **0.145** | 0.152 | 0.084 | 0.084 | 0.960 | 0.440 | 0.215 | 0.064 |
| $\log(OR^{NDE})$ | **0.100** | 0.107 | 0.085 | 0.085 | 0.955 | 0.237 | 0.082 | 0.015 |
| $\log(OR^{NIE})$ | **0.045** | 0.045 | 0.021 | 0.020 | 0.948 | 0.615 | 0.252 | 0.012 |
| $\log(OR^{CDE})$ | **0.100** | 0.107 | 0.085 | 0.085 | 0.955 | 0.237 | 0.082 | 0.015 |

### TABLE S7: CONTINUOUS MEDIATOR WITH INTERACTION EFFECT, DISEASE PREVALENCE OF 0.20

| | | | | | | Power at $\alpha =$ | | |
|---|---|---|---|---|---|---|---|---|
| | True Value | Mean Value | Mean SE | Emp. SD | 95% CI Coverage | 0.05 | 0.01 | 0.001 |
| **Likelihood** | | | | | | | | |
| $\log(OR^{TE})$ | **0.145** | 0.135 | 0.084 | 0.082 | 0.953 | 0.343 | 0.163 | 0.040 |
| $\log(OR^{NDE})$ | **0.100** | 0.098 | 0.085 | 0.083 | 0.952 | 0.207 | 0.065 | 0.014 |
| $\log(OR^{NIE})$ | **0.045** | 0.037 | 0.021 | 0.020 | 0.951 | 0.484 | 0.208 | 0.030 |
| $\log(OR^{CDE})$ | **0.077** | 0.077 | 0.087 | 0.085 | 0.948 | 0.142 | 0.039 | 0.008 |
| **VW** | | | | | | | | |
| $\log(OR^{TE})$ | **0.145** | 0.137 | 0.089 | 0.083 | 0.956 | 0.310 | 0.132 | 0.028 |
| $\log(OR^{NDE})$ | **0.100** | 0.100 | 0.086 | 0.084 | 0.951 | 0.210 | 0.064 | 0.013 |
| $\log(OR^{NIE})$ | **0.045** | 0.038 | 0.027 | 0.022 | 0.965 | 0.162 | 0.006 | 0.000 |
| $\log(OR^{CDE})$ | **0.077** | 0.077 | 0.087 | 0.086 | 0.951 | 0.145 | 0.038 | 0.005 |
| **VW-IPW** | | | | | | | | |
| $\log(OR^{TE})$ | **0.145** | 0.144 | 0.089 | 0.086 | 0.957 | 0.340 | 0.154 | 0.031 |
| $\log(OR^{NDE})$ | **0.100** | 0.102 | 0.086 | 0.085 | 0.948 | 0.211 | 0.068 | 0.013 |
| $\log(OR^{NIE})$ | **0.045** | 0.042 | 0.027 | 0.024 | 0.969 | 0.317 | 0.053 | 0.000 |
| $\log(OR^{CDE})$ | **0.077** | 0.077 | 0.087 | 0.086 | 0.951 | 0.145 | 0.038 | 0.005 |

#### TABLE S8: TYPE-I ERROR RATES FOR TESTING DIRECT AND INDIRECT EFFECTS

| Mediator | | Direct Effect ($\gamma_A = 0$) | | Indirect Effect ($\gamma_M = 0$) | |
|---|---|---|---|---|---|
| | | $\alpha = 0.05$ | $\alpha = 0.01$ | $\alpha = 0.05$ | $\alpha = 0.01$ |
| Continuous | | | | | |
| | Likelihood | 0.0492 | 0.0104 | 0.0339 | 0.0042 |
| | VW | 0.0493 | 0.0104 | 0.0194 | 0.0011 |
| | VW-IPW | 0.0492 | 0.0104 | 0.0202 | 0.0011 |
| Binary | | | | | |
| | Likelihood | 0.0471 | 0.0104 | 0.0319 | 0.0035 |
| | VW | 0.0471 | 0.0104 | 0.0195 | 0.0015 |
| | VW-IPW | 0.0471 | 0.0104 | 0.0205 | 0.0015 |

### TABLE S9: BINARY MEDIATOR WITH NO INTERACTION

| | True Value | Mean Value | Mean SE | SD |
|---|---|---|---|---|
| **Likelihood** | | | | |
| $\gamma_A$ | **0.100** | 0.097 | 0.085 | 0.085 |
| $\gamma_M$ | **0.420** | 0.428 | 0.169 | 0.179 |
| $\gamma_C$ | **0.100** | 0.111 | 0.166 | 0.163 |
| $\beta_0$ | **-0.310** | -0.326 | 0.147 | 0.151 |
| $\beta_A$ | **0.430** | 0.434 | 0.088 | 0.090 |
| $\beta_C$ | **0.100** | 0.087 | 0.169 | 0.171 |
| **VW** | | | | |
| $\gamma_A$ | **0.100** | 0.097 | 0.085 | 0.085 |
| $\gamma_M$ | **0.420** | 0.428 | 0.169 | 0.179 |
| $\gamma_C$ | **0.100** | 0.111 | 0.166 | 0.163 |
| $\beta_0$ | **-0.310** | -0.325 | 0.170 | 0.171 |
| $\beta_A$ | **0.430** | 0.432 | 0.126 | 0.130 |
| $\beta_C$ | **0.100** | 0.084 | 0.240 | 0.234 |
| **VW-IPW** | | | | |
| $\gamma_A$ | **0.100** | 0.097 | 0.085 | 0.085 |
| $\gamma_M$ | **0.420** | 0.428 | 0.169 | 0.179 |
| $\gamma_C$ | **0.100** | 0.111 | 0.166 | 0.163 |
| $\beta_0$ | **-0.310** | -0.313 | 0.166 | 0.166 |
| $\beta_A$ | **0.430** | 0.433 | 0.123 | 0.126 |
| $\beta_C$ | **0.100** | 0.085 | 0.235 | 0.227 |

### TABLE S10: COVARIANCE ESTIMATES FOR BINARY MEDIATOR (NO INTERACTION)

| **Likelihood** | | | | | | |
|---|---|---|---|---|---|---|
| | $\gamma_A$ | $\gamma_M$ | $\gamma_C$ | $\beta_0$ | $\beta_A$ | $\beta_C$ |
| $\gamma_A$ | 0.0072 | -0.0029 | 0.0001 | 0.0015 | -0.0007 | 0.0001 |
| $\gamma_M$ | -0.0029 | 0.0286 | -0.0005 | -0.0139 | -0.0004 | -0.0007 |
| $\gamma_C$ | 0.0001 | -0.0005 | 0.0274 | 0.0017 | 0 | -0.0029 |
| $\beta_0$ | 0.0015 | -0.0139 | 0.0017 | 0.0215 | -0.0004 | -0.0144 |
| $\beta_A$ | -0.0007 | -0.0004 | 0 | -0.0004 | 0.0078 | 0.0001 |
| $\beta_C$ | 0.0001 | -0.0007 | -0.0029 | -0.0144 | 0.0001 | 0.0286 |
| **VW** | | | | | | |
| $\gamma_A$ | 0.0072 | -0.0029 | 0.0001 | 0 | 0 | 0 |
| $\gamma_M$ | -0.0029 | 0.0286 | -0.0005 | 0 | 0 | 0 |
| $\gamma_C$ | 0.0001 | -0.0005 | 0.0274 | 0 | 0 | 0 |
| $\beta_0$ | 0 | 0 | 0 | 0.029 | -0.0009 | -0.0289 |
| $\beta_A$ | 0 | 0 | 0 | -0.0009 | 0.0158 | 0.0003 |
| $\beta_C$ | 0 | 0 | 0 | -0.0289 | 0.0003 | 0.0577 |
| **VW-IPW** | | | | | | |
| $\gamma_A$ | 0.0072 | -0.0029 | 0.0001 | 0 | 0 | 0 |
| $\gamma_M$ | -0.0029 | 0.0286 | -0.0005 | 0 | 0 | 0 |
| $\gamma_C$ | 0.0001 | -0.0005 | 0.0274 | 0 | 0 | 0 |
| $\beta_0$ | 0 | 0 | 0 | 0.0277 | -0.0009 | -0.0277 |
| $\beta_A$ | 0 | 0 | 0 | -0.0009 | 0.0153 | 0.0002 |
| $\beta_C$ | 0 | 0 | 0 | -0.0277 | 0.0002 | 0.0551 |

### TABLE S11: BINARY MEDIATOR WITH INTERACTION

| | | | | | | | Power at $\alpha$ = | | |
|---|---|---|---|---|---|---|---|---|---|
| | | True Value | Mean Value | Mean SE | SD | 95% CI Coverage | 0.05 | 0.01 | 0.001 |
| **Likelihood** | | | | | | | | | |
| $\log(OR^{TE})$ | | **0.145** | 0.148 | 0.084 | 0.085 | 0.946 | 0.427 | 0.214 | 0.053 |
| $\log(OR^{NDE})$ | | **0.100** | 0.107 | 0.085 | 0.084 | 0.953 | 0.243 | 0.080 | 0.023 |
| $\log(OR^{NIE})$ | | **0.045** | 0.041 | 0.023 | 0.023 | 0.969 | 0.501 | 0.218 | 0.029 |
| $\log(OR^{CDE})$ | | **0.049** | 0.055 | 0.117 | 0.117 | 0.953 | 0.077 | 0.018 | 0.002 |
| **VW** | | | | | | | | | |
| $\log(OR^{TE})$ | | **0.145** | 0.148 | 0.088 | 0.085 | 0.961 | 0.392 | 0.168 | 0.036 |
| $\log(OR^{NDE})$ | | **0.100** | 0.107 | 0.086 | 0.084 | 0.951 | 0.233 | 0.079 | 0.021 |
| $\log(OR^{NIE})$ | | **0.045** | 0.041 | 0.029 | 0.023 | 0.986 | 0.193 | 0.020 | 0.000 |
| $\log(OR^{CDE})$ | | **0.049** | 0.055 | 0.118 | 0.118 | 0.951 | 0.075 | 0.017 | 0.002 |
| **VW-IPW** | | | | | | | | | |
| $\log(OR^{TE})$ | | **0.145** | 0.149 | 0.088 | 0.085 | 0.959 | 0.392 | 0.172 | 0.037 |
| $\log(OR^{NDE})$ | | **0.100** | 0.108 | 0.086 | 0.084 | 0.951 | 0.235 | 0.079 | 0.021 |
| $\log(OR^{NIE})$ | | **0.045** | 0.041 | 0.029 | 0.023 | 0.986 | 0.224 | 0.020 | 0.000 |
| $\log(OR^{CDE})$ | | **0.049** | 0.055 | 0.118 | 0.118 | 0.951 | 0.075 | 0.017 | 0.002 |

### TABLE S12: BINARY MEDIATOR WITH INTERACTION

| | True Value | Mean Value | Mean SE | SD |
|---|---|---|---|---|
| **Likelihood** | | | | |
| $\gamma_A$ | **0.049** | 0.056 | 0.117 | 0.117 |
| $\gamma_M$ | **0.320** | 0.321 | 0.170 | 0.167 |
| $\gamma_{AM}$ | **0.100** | 0.093 | 0.169 | 0.173 |
| $\gamma_C$ | **0.100** | 0.099 | 0.165 | 0.166 |
| $\beta_0$ | **-0.310** | -0.320 | 0.147 | 0.145 |
| $\beta_A$ | **0.430** | 0.434 | 0.123 | 0.122 |
| $\beta_C$ | **0.100** | 0.094 | 0.170 | 0.171 |
| **VW** | | | | |
| $\gamma_A$ | **0.049** | 0.057 | 0.118 | 0.118 |
| $\gamma_M$ | **0.320** | 0.321 | 0.170 | 0.167 |
| $\gamma_{AM}$ | **0.100** | 0.093 | 0.170 | 0.175 |
| $\gamma_C$ | **0.100** | 0.099 | 0.165 | 0.166 |
| $\beta_0$ | **-0.310** | -0.321 | 0.170 | 0.173 |
| $\beta_A$ | **0.430** | 0.436 | 0.126 | 0.125 |
| $\beta_C$ | **0.100** | 0.095 | 0.240 | 0.253 |
| **VW-IPW** | | | | |
| $\gamma_A$ | **0.049** | 0.057 | 0.118 | 0.118 |
| $\gamma_M$ | **0.320** | 0.321 | 0.170 | 0.167 |
| $\gamma_{AM}$ | **0.100** | 0.093 | 0.170 | 0.175 |
| $\gamma_C$ | **0.100** | 0.099 | 0.165 | 0.166 |
| $\beta_0$ | **-0.310** | -0.312 | 0.167 | 0.167 |
| $\beta_A$ | **0.430** | 0.439 | 0.124 | 0.121 |
| $\beta_C$ | **0.100** | 0.096 | 0.235 | 0.244 |

### TABLE S13: COVARIANCE ESTIMATES FOR BINARY MEDIATOR (WITH INTERACTION)

| **Likelihood** | | | | | | | |
|---|---|---|---|---|---|---|---|
| | $\gamma_A$ | $\gamma_M$ | $\gamma_{AM}$ | $\gamma_C$ | $\beta_0$ | $\beta_A$ | $\beta_C$ |
| $\gamma_A$ | 0.0138 | -0.0022 | -0.0138 | 0.0001 | 0.0011 | 0.0064 | 0.0001 |
| $\gamma_M$ | -0.0022 | 0.029 | -0.0022 | -0.0006 | -0.0141 | 0.0006 | -0.0007 |
| $\gamma_{AM}$ | -0.0138 | -0.0022 | 0.0287 | 0 | 0.0007 | -0.0143 | 0 |
| $\gamma_C$ | 0.0001 | -0.0006 | 0 | 0.0273 | 0.0014 | 0 | -0.0022 |
| $\beta_0$ | 0.0011 | -0.0141 | 0.0007 | 0.0014 | 0.0216 | -0.0008 | -0.0144 |
| $\beta_A$ | 0.0064 | 0.0006 | -0.0143 | 0 | -0.0008 | 0.0151 | 0.0002 |
| $\beta_C$ | 0.0001 | -0.0007 | 0 | -0.0022 | -0.0144 | 0.0002 | 0.0288 |
| **VW** | | | | | | | |
| $\gamma_A$ | 0.0139 | -0.0022 | -0.0139 | 0.0001 | 0 | 0 | 0 |
| $\gamma_M$ | -0.0022 | 0.029 | -0.0022 | -0.0006 | 0 | 0 | 0 |
| $\gamma_{AM}$ | -0.0139 | -0.0022 | 0.0291 | 0 | 0 | 0 | 0 |
| $\gamma_C$ | 0.0001 | -0.0006 | 0 | 0.0274 | 0 | 0 | 0 |
| $\beta_0$ | 0 | 0 | 0 | 0 | 0.029 | -0.0009 | -0.0289 |
| $\beta_A$ | 0 | 0 | 0 | 0 | -0.0009 | 0.0159 | 0.0003 |
| $\beta_C$ | 0 | 0 | 0 | 0 | -0.0289 | 0.0003 | 0.0577 |
| **VW-IPW** | | | | | | | |
| $\gamma_A$ | 0.0139 | -0.0022 | -0.0139 | 0.0001 | 0 | 0 | 0 |
| $\gamma_M$ | -0.0022 | 0.029 | -0.0022 | -0.0006 | 0 | 0 | 0 |
| $\gamma_{AM}$ | -0.0139 | -0.0022 | 0.0291 | 0 | 0 | 0 | 0 |
| $\gamma_C$ | 0.0001 | -0.0006 | 0 | 0.0274 | 0 | 0 | 0 |
| $\beta_0$ | 0 | 0 | 0 | 0 | 0.0278 | -0.0009 | -0.0278 |
| $\beta_A$ | 0 | 0 | 0 | 0 | -0.0009 | 0.0153 | 0.0004 |
| $\beta_C$ | 0 | 0 | 0 | 0 | -0.0278 | 0.0004 | 0.0552 |

**TABLE S14: BINARY MEDIATOR, DISEASE PREVALENCE OF 0.20**

| | True Value | Mean Value | Mean SE | Emp. SD | 95% CI Coverage | Power at $\alpha =$ 0.05 | 0.01 | 0.001 |
|---|---|---|---|---|---|---|---|---|
| **Likelihood** | | | | | | | | |
| $\log(OR^{TE})$ | **0.145** | 0.147 | 0.083 | 0.086 | 0.947 | 0.433 | 0.224 | 0.060 |
| $\log(OR^{NDE})$ | **0.100** | 0.101 | 0.085 | 0.088 | 0.940 | 0.228 | 0.079 | 0.021 |
| $\log(OR^{NIE})$ | **0.045** | 0.046 | 0.020 | 0.020 | 0.945 | 0.657 | 0.310 | 0.037 |
| $\log(OR^{CDE})$ | **0.100** | 0.101 | 0.085 | 0.088 | 0.940 | 0.229 | 0.079 | 0.021 |
| **VW** | | | | | | | | |
| $\log(OR^{TE})$ | **0.145** | 0.148 | 0.085 | 0.087 | 0.945 | 0.416 | 0.204 | 0.054 |
| $\log(OR^{NDE})$ | **0.100** | 0.101 | 0.085 | 0.088 | 0.940 | 0.229 | 0.079 | 0.021 |
| $\log(OR^{NIE})$ | **0.045** | 0.046 | 0.023 | 0.023 | 0.931 | 0.483 | 0.129 | 0.007 |
| $\log(OR^{CDE})$ | **0.100** | 0.101 | 0.085 | 0.088 | 0.940 | 0.229 | 0.079 | 0.021 |
| **VW-IPW** | | | | | | | | |
| $\log(OR^{TE})$ | **0.145** | 0.148 | 0.084 | 0.087 | 0.945 | 0.420 | 0.208 | 0.055 |
| $\log(OR^{NDE})$ | **0.100** | 0.101 | 0.085 | 0.088 | 0.940 | 0.229 | 0.079 | 0.021 |
| $\log(OR^{NIE})$ | **0.045** | 0.047 | 0.021 | 0.022 | 0.942 | 0.609 | 0.269 | 0.030 |
| $\log(OR^{CDE})$ | **0.100** | 0.101 | 0.085 | 0.088 | 0.940 | 0.229 | 0.079 | 0.021 |

#### TABLE S15: BINARY MEDIATOR WITH INTERACTION EFFECT, DISEASE PREVALENCE OF 0.20

| | True Value | Mean Value | Mean SE | Emp. SD | 95% CI Coverage | Power at $\alpha =$ 0.05 | 0.01 | 0.001 |
|---|---|---|---|---|---|---|---|---|
| **Likelihood** | | | | | | | | |
| $\log(OR^{TE})$ | **0.145** | 0.145 | 0.084 | 0.086 | 0.951 | 0.399 | 0.197 | 0.050 |
| $\log(OR^{NDE})$ | **0.100** | 0.105 | 0.085 | 0.086 | 0.950 | 0.233 | 0.083 | 0.017 |
| $\log(OR^{NIE})$ | **0.045** | 0.040 | 0.022 | 0.022 | 0.950 | 0.509 | 0.214 | 0.022 |
| $\log(OR^{CDE})$ | **0.049** | 0.047 | 0.115 | 0.115 | 0.947 | 0.071 | 0.018 | 0.001 |
| **VW** | | | | | | | | |
| $\log(OR^{TE})$ | **0.145** | 0.146 | 0.089 | 0.087 | 0.959 | 0.367 | 0.162 | 0.034 |
| $\log(OR^{NDE})$ | **0.100** | 0.105 | 0.086 | 0.087 | 0.955 | 0.230 | 0.080 | 0.017 |
| $\log(OR^{NIE})$ | **0.045** | 0.040 | 0.028 | 0.023 | 0.973 | 0.178 | 0.008 | 0.000 |
| $\log(OR^{CDE})$ | **0.049** | 0.047 | 0.115 | 0.115 | 0.949 | 0.069 | 0.016 | 0.001 |
| **VW-IPW** | | | | | | | | |
| $\log(OR^{TE})$ | **0.145** | 0.151 | 0.090 | 0.089 | 0.956 | 0.398 | 0.184 | 0.037 |
| $\log(OR^{NDE})$ | **0.100** | 0.107 | 0.086 | 0.087 | 0.950 | 0.237 | 0.083 | 0.018 |
| $\log(OR^{NIE})$ | **0.045** | 0.044 | 0.027 | 0.024 | 0.970 | 0.332 | 0.073 | 0.003 |
| $\log(OR^{CDE})$ | **0.049** | 0.047 | 0.115 | 0.115 | 0.949 | 0.069 | 0.016 | 0.001 |

### TABLE S16: CONTINUOUS MEDIATOR WITH NO INTERACTION: NAÏVE ANALYSIS

| | True Value | Mean Value | Mean SE | SD |
|---|---|---|---|---|
| **Likelihood** | | | | |
| $\gamma_A$ | **0.100** | 0.099 | 0.085 | 0.083 |
| $\gamma_M$ | **0.300** | 0.304 | 0.118 | 0.114 |
| $\gamma_C$ | **0.050** | 0.052 | 0.166 | 0.167 |
| $\beta_0$ | **0.100** | 0.095 | 0.050 | 0.049 |
| $\beta_A$ | **0.150** | 0.149 | 0.029 | 0.028 |
| $\beta_C$ | **0.050** | 0.048 | 0.058 | 0.057 |
| $\sigma^2$ | **0.500** | 0.498 | 0.029 | 0.028 |
| **VW** | | | | |
| $\gamma_A$ | **0.100** | 0.099 | 0.085 | 0.083 |
| $\gamma_M$ | **0.300** | 0.304 | 0.118 | 0.114 |
| $\gamma_C$ | **0.050** | 0.052 | 0.166 | 0.167 |
| $\beta_0$ | **0.100** | 0.094 | 0.058 | 0.056 |
| $\beta_A$ | **0.150** | 0.149 | 0.041 | 0.040 |
| $\beta_C$ | **0.050** | 0.050 | 0.082 | 0.080 |
| $\sigma^2$ | **0.500** | 0.496 | 0.041 | 0.042 |
| **VW-NAIVE** | | | | |
| $\gamma_A$ | **0.100** | 0.099 | 0.085 | 0.083 |
| $\gamma_M$ | **0.300** | 0.304 | 0.118 | 0.114 |
| $\gamma_C$ | **0.050** | 0.052 | 0.166 | 0.167 |
| $\beta_0$ | **0.100** | 0.169 | 0.041 | 0.041 |
| $\beta_A$ | **0.150** | 0.155 | 0.029 | 0.028 |
| $\beta_C$ | **0.050** | 0.051 | 0.058 | 0.057 |
| $\sigma^2$ | **0.500** | 0.507 | 0.029 | 0.028 |

Assumes simulation setup for continuous mediator in Table 1

### TABLE S17: CONTINUOUS MEDIATOR WITH NO INTERACTION, NAÏVE ANALYSIS

| | True Value | Mean Value | Mean SE | SD | 95% CI Coverage | Power at $\alpha =$ 0.05 | 0.01 | 0.001 |
|---|---|---|---|---|---|---|---|---|
| **Likelihood** | | | | | | | | |
| $\log(OR^{TE})$ | **0.145** | 0.144 | 0.083 | 0.081 | 0.961 | 0.408 | 0.198 | 0.055 |
| $\log(OR^{NDE})$ | **0.100** | 0.099 | 0.085 | 0.083 | 0.958 | 0.199 | 0.072 | 0.014 |
| $\log(OR^{NIE})$ | **0.045** | 0.045 | 0.019 | 0.019 | 0.956 | 0.691 | 0.362 | 0.058 |
| $\log(OR^{CDE})$ | **0.100** | 0.099 | 0.085 | 0.083 | 0.958 | 0.199 | 0.072 | 0.014 |
| **VW** | | | | | | | | |
| $\log(OR^{TE})$ | **0.145** | 0.144 | 0.084 | 0.081 | 0.964 | 0.397 | 0.182 | 0.046 |
| $\log(OR^{NDE})$ | **0.100** | 0.099 | 0.085 | 0.083 | 0.958 | 0.199 | 0.072 | 0.014 |
| $\log(OR^{NIE})$ | **0.045** | 0.045 | 0.022 | 0.021 | 0.940 | 0.504 | 0.131 | 0.009 |
| $\log(OR^{CDE})$ | **0.100** | 0.099 | 0.085 | 0.083 | 0.958 | 0.199 | 0.072 | 0.014 |
| **VW-NAIVE** | | | | | | | | |
| $\log(OR^{TE})$ | **0.145** | 0.144 | 0.084 | 0.081 | 0.963 | 0.399 | 0.190 | 0.051 |
| $\log(OR^{NDE})$ | **0.100** | 0.099 | 0.085 | 0.083 | 0.958 | 0.199 | 0.072 | 0.014 |
| $\log(OR^{NIE})$ | **0.045** | 0.045 | 0.020 | 0.021 | 0.924 | 0.640 | 0.284 | 0.042 |
| $\log(OR^{CDE})$ | **0.100** | 0.099 | 0.085 | 0.083 | 0.958 | 0.199 | 0.072 | 0.014 |

Assumes simulation setup for continuous mediator in Table 1

### TABLE S18: CONTINUOUS MEDIATOR WITH NO INTERACTION: NAÏVE ANALYSIS, RARER OUTCOME

| | True Value | Mean Value | Mean SE | SD |
|---|---|---|---|---|
| **Likelihood** | | | | |
| $\gamma_A$ | **0.400** | 0.404 | 0.090 | 0.091 |
| $\gamma_M$ | **0.600** | 0.597 | 0.125 | 0.123 |
| $\gamma_C$ | **0.050** | 0.060 | 0.173 | 0.169 |
| $\beta_0$ | **0.100** | 0.100 | 0.050 | 0.049 |
| $\beta_A$ | **0.150** | 0.151 | 0.029 | 0.029 |
| $\beta_C$ | **0.050** | 0.051 | 0.058 | 0.057 |
| $\sigma^2$ | **0.500** | 0.496 | 0.029 | 0.028 |
| **VW** | | | | |
| $\gamma_A$ | **0.400** | 0.404 | 0.090 | 0.091 |
| $\gamma_M$ | **0.600** | 0.597 | 0.125 | 0.123 |
| $\gamma_C$ | **0.050** | 0.060 | 0.173 | 0.169 |
| $\beta_0$ | **0.100** | 0.101 | 0.057 | 0.057 |
| $\beta_A$ | **0.150** | 0.153 | 0.041 | 0.043 |
| $\beta_C$ | **0.050** | 0.051 | 0.081 | 0.083 |
| $\sigma^2$ | **0.500** | 0.496 | 0.040 | 0.039 |
| **VW-NAIVE** | | | | |
| $\gamma_A$ | **0.400** | 0.404 | 0.090 | 0.091 |
| $\gamma_M$ | **0.600** | 0.597 | 0.125 | 0.123 |
| $\gamma_C$ | **0.050** | 0.060 | 0.173 | 0.169 |
| $\beta_0$ | **0.100** | 0.237 | 0.043 | 0.041 |
| $\beta_A$ | **0.150** | 0.195 | 0.029 | 0.029 |
| $\beta_C$ | **0.050** | 0.058 | 0.059 | 0.058 |
| $\sigma^2$ | **0.500** | 0.520 | 0.029 | 0.030 |

Assumes disease prevalence of 0.003, $\gamma_A = 0.4$, and $\gamma_M = 0.6$. All other simulation parameters are those reported for continuous mediator in Table 1.

### TABLE S19: CONTINUOUS MEDIATOR WITH NO INTERACTION, NAÏVE ANALYSIS, RARER OUTCOME

| | True Value | Mean Value | Mean SE | SD | 95% CI Coverage | Power at $\alpha =$ 0.05 | 0.01 | 0.001 |
|---|---|---|---|---|---|---|---|---|
| **Likelihood** | | | | | | | | |
| $\log(OR^{TE})$ | **0.490** | 0.494 | 0.087 | 0.088 | 0.951 | 1.000 | 1.000 | 0.996 |
| $\log(OR^{NDE})$ | **0.400** | 0.404 | 0.090 | 0.091 | 0.949 | 0.997 | 0.973 | 0.907 |
| $\log(OR^{NIE})$ | **0.090** | 0.089 | 0.023 | 0.022 | 0.950 | 0.998 | 0.991 | 0.864 |
| $\log(OR^{CDE})$ | **0.400** | 0.404 | 0.090 | 0.091 | 0.949 | 0.997 | 0.973 | 0.907 |
| **VW** | | | | | | | | |
| $\log(OR^{TE})$ | **0.490** | 0.494 | 0.092 | 0.089 | 0.952 | 1.000 | 1.000 | 0.992 |
| $\log(OR^{NDE})$ | **0.400** | 0.404 | 0.090 | 0.091 | 0.949 | 0.997 | 0.973 | 0.910 |
| $\log(OR^{NIE})$ | **0.090** | 0.090 | 0.031 | 0.029 | 0.957 | 0.930 | 0.694 | 0.174 |
| $\log(OR^{CDE})$ | **0.400** | 0.404 | 0.090 | 0.091 | 0.949 | 0.997 | 0.973 | 0.910 |
| **VW-NAIVE** | | | | | | | | |
| $\log(OR^{TE})$ | **0.490** | 0.494 | 0.091 | 0.089 | 0.948 | 1.000 | 1.000 | 0.994 |
| $\log(OR^{NDE})$ | **0.400** | 0.404 | 0.090 | 0.091 | 0.949 | 0.997 | 0.973 | 0.910 |
| $\log(OR^{NIE})$ | **0.090** | 0.090 | 0.026 | 0.029 | 0.901 | 0.985 | 0.903 | 0.591 |
| $\log(OR^{CDE})$ | **0.400** | 0.404 | 0.090 | 0.091 | 0.949 | 0.997 | 0.973 | 0.910 |

Assumes disease prevalence of 0.003, $\gamma_A = 0.4$, and $\gamma_M = 0.6$. All other simulation parameters are those reported for continuous mediator in Table 1.

### TABLE S20: PARAMETER ESTIMATES FOR MEDIATION ANALYSIS OF *rs12914385* AND LUNG CANCER

| | Likelihood | | | | VW | | |
|---|---|---|---|---|---|---|---|
| | Estimate | SE | P-value | | Estimate | SE | P-value |
| $\gamma_A$ | 0.2871 | 0.0427 | <0.0001 | | 0.2876 | 0.0427 | <0.0001 |
| $\gamma_M$ | 1.623 | 0.0636 | <0.0001 | | 1.62412 | 0.06369 | <0.0001 |
| $\gamma_{C,gender}$ | 0.3003 | 0.0712 | <0.0001 | | 0.3025 | 0.0712 | <0.0001 |
| $\gamma_{C,PC1}$ | 6.8498 | 2.3088 | 0.0030 | | 7.0410 | 2.3153 | 0.0024 |
| $\gamma_{C,PC2}$ | -2.4621 | 2.2704 | 0.2782 | | -2.3686 | 2.2711 | 0.29706 |
| $\gamma_{C,PC3}$ | 2.4194 | 2.2380 | 0.2797 | | 2.4739 | 2.2397 | 0.2693 |
| $\beta_0$ | 0.8486 | 0.1045 | <0.0001 | | 0.4857 | 0.1438 | 0.0007 |
| $\beta_A$ | 0.0889 | 0.0442 | 0.0445 | | 0.0394 | 0.0609 | 0.5173 |
| $\beta_{C,gender}$ | -1.2681 | 0.0741 | <0.0001 | | -0.9424 | 0.1042 | <0.0001 |
| $\beta_{C,PC1}$ | -25.4531 | 2.4406 | <0.0001 | | -38.8204 | 3.2266 | <0.0001 |
| $\beta_{C,PC2}$ | 17.1941 | 2.3453 | <0.0001 | | 22.8812 | 3.3611 | <0.0001 |
| $\beta_{C,PC3}$ | 0.4251 | 2.3144 | 0.8543 | | 1.2488 | 3.1396 | 0.6908 |

| | VW-IPW | | |
|---|---|---|---|
| | Estimate | SE | P-value |
| $\gamma_A$ | 0.2876 | 0.0427 | <0.0001 |
| $\gamma_M$ | 1.62412 | 0.06369 | <0.0001 |
| $\gamma_{C,gender}$ | 0.3025 | 0.0712 | <0.0001 |
| $\gamma_{C,PC1}$ | 7.0410 | 2.3153 | 0.0024 |
| $\gamma_{C,PC2}$ | -2.3686 | 2.2711 | 0.29706 |
| $\gamma_{C,PC3}$ | 2.4739 | 2.2397 | 0.2693 |
| $\beta_0$ | 0.58943 | 0.1019 | <0.0001 |
| $\beta_A$ | 0.07451 | 0.0423 | 0.0831 |
| $\beta_{C,gender}$ | -0.9652 | 0.0737 | <0.0001 |
| $\beta_{C,PC1}$ | -36.2963 | 2.3016 | <0.0001 |
| $\beta_{C,PC2}$ | 21.7720 | 2.3600 | <0.0001 |
| $\beta_{C,PC3}$ | 1.3304 | 2.2256 | 0.5500 |

### SUPPLEMENTAL REFERENCES

1. Valeri L, VanderWeele TJ. Mediation analysis allowing for exposure–mediator interactions and causal interpretation: theoretical assumptions and implementation with SAS and SPSS macros. *Psychological methods*. 2013;18(2):137.
